# 5-ethynyluridine perturbs nuclear RNA metabolism to promote the nuclear accumulation of TDP-43 and other RNA binding proteins

**DOI:** 10.1101/2025.04.02.646885

**Authors:** Lindsey R. Hayes, Benjamin Zaepfel, Lauren Duan, Anne C. Starner, Mason D. Bartels, Rebekah L. Rothacher, Sophie Martin, Rachel French, Zhe Zhang, Irika R. Sinha, Jonathan P. Ling, Shuying Sun, Yuna M. Ayala, Jeff Coller, Eric L. Van Nostrand, Liliana Florea, Petr Kalab

## Abstract

TDP-43, an essential nucleic acid binding protein and splicing regulator, is broadly disrupted in neurodegeneration. TDP-43 nuclear localization and function depend on the abundance of its nuclear RNA targets and its recruitment into large ribonucleoprotein complexes, which restricts TDP-43 nuclear efflux. To further investigate the interplay between TDP-43 and nascent RNAs, we aimed to employ 5-ethynyluridine (5EU), a widely used uridine analog for ‘click chemistry’ labeling of newly transcribed RNAs. Surprisingly, 5EU induced the nuclear accumulation of TDP-43 and other RNA-binding proteins and attenuated TDP-43 mislocalization caused by disruption of the nuclear transport apparatus. RNA FISH demonstrated 5EU-induced nuclear accumulation of polyadenylated and GU-repeat-rich RNAs, suggesting increased retention of both processed and intronic RNAs. TDP-43 eCLIP confirmed that 5EU preserved TDP-43 binding at predominantly GU-rich intronic sites. RNAseq revealed significant 5EU-induced changes in alternative splicing, accompanied by an overall reduction in splicing diversity, without any major changes in RNA stability or TDP-43 splicing regulatory function. These data suggest that 5EU may impede RNA splicing efficiency and subsequent nuclear RNA processing and export. Our findings have important implications for studies utilizing 5EU and offer unexpected confirmation that the accumulation of endogenous nuclear RNAs promotes TDP-43 nuclear localization.

## INTRODUCTION

The molecular mechanisms regulating the localization and function of the essential nucleic acid binding protein, TAR-DNA binding protein (TDP-43), are of major interest in neurodegenerative diseases, particularly the TDP-43 proteinopathies,^1^ a group of age-related disorders including amyotrophic lateral sclerosis (ALS), frontotemporal dementia (FTD), and limbic-predominant age-related TDP-43 encephalopathy (LATE). Loss of the nuclear splicing regulatory function of TDP-43 induces cell-specific alternative splicing,^2,3^ failure of cryptic exon repression,^4–6^ and alternative polyadenylation,^7–9^ that has been shown to disrupt numerous targets, including neuronal genes critical for axonal and synaptic function.^10–13^ Neuropathology shows TDP-43 nuclear clearance and mislocalization to the cytoplasm in neurons and glia in affected regions of the central nervous system (CNS), accompanied by the formation of cytoplasmic aggregates.^14–16^ Disruption of TDP-43 nucleocytoplasmic transport and perturbation of TDP-43 solubility, through disruption of TDP-43 liquid-liquid phase separation, autoregulation, and turnover, have all been implicated in the TDP-43 pathogenic cascade.^17,18^ Alleviating disruptions of these pathways in disease models supports TDP-43 as a therapeutic target, though findings still await translation to patients.^19^ Improved understanding of the regulation of TDP-43 homeostasis is needed to expand our molecular toolbox for experimental and therapeutic modulation of TDP-43.

We and others have shown that the normal predominantly nuclear localization of 43-kD full-length TDP-43 results from facilitated nuclear import and passive exit through the nuclear pore complex (NPC).^20–24^ Binding of importins alpha and beta to the TDP-43 N-terminal nuclear localization signal (NLS) facilitates its nuclear entry, driven by the nucleocytoplasmic gradient of RanGTP. Importin beta also binds TDP-43 in its C-terminal intrinsically disordered domain, contributing to non-NLS-mediated nuclear import.^25^ TDP-43 nuclear retention, in turn, depends on TDP-43 nuclear RNA binding and incorporation into high molecular weight ribonucleoprotein (RNP) complexes,^21,26–28^ limiting its passive exit through NPC channels, which increasingly restrict the containing RNP complexes far exceed the passive limits of the NPC.^26,30^ Disruption of TDP-43 RNA binding or depletion of nuclear RNAs available for TDP-43 to bind markedly disrupts its nuclear localization.^21^ Disruption of TDP-43 phase separation and N-terminal oligomerization, which are critical for TDP-43 homo- and hetero-oligomerization within RNP complexes, also perturbs its nuclear localization.^26,28^ Conversely, we demonstrated that increasing the availability of TDP-43 nuclear RNA binding sites, particularly its preferred GU-rich motifs found within introns, promotes TDP-43 nuclear accumulation.^21,27^ Small molecule inhibitors of splicing,^21^ acute blockade of nuclear RNA export,^21^ and introduction of synthetic multivalent GU-rich oligonucleotides^27^ all promote TDP-43 nuclear accumulation, supporting a model in which the localization of TDP-43 strongly depends on the subcellular localization of its RNA targets.

To further investigate the relationship between TDP-43 and RNA localization within cells, we set out to use 5-ethynyluridine (5EU) to mark newly transcribed RNAs via ‘click chemistry’ labeling.^31^ 5EU is a well-established and widely used RNA metabolic label that is incorporated into newly transcribed RNAs (by RNA Pol I, II, and III) for analysis of RNA transcription, transport, and turnover.^32^ The broad utility of 5EU for RNA imaging and sequencing studies has led to numerous advances in RNA biology.^33^ Unexpectedly, we found that 5EU induced the time- and dose-dependent nuclear accumulation of TDP-43 and related RNA binding proteins (RBPs) in cell lines and neurons, and attenuated TDP-43 mislocalization after transcriptional blockade or disruption of the nuclear transport apparatus. Further mechanistic studies demonstrated 5EU-induced accumulation of nuclear RNAs, including polyadenylated and GU-rich nuclear RNAs. TDP-43 eCLIP showed that 5EU preserves TDP-43 binding to GU-rich intronic sites that are normally depleted by transcriptional blockade. Transcriptome-wide analysis of RNA decay did not show major effects of 5EU on RNA stability. However, widespread changes in gene expression were seen in 5EU-treated cells, including alterations in RNA splicing and metabolic pathways, as well as marked changes in alternative splicing with an overall decrease in splicing diversity. Our findings suggest that 5EU-induced changes in splicing and subsequent nuclear RNA metabolism and export promote the accumulation of nuclear RNAs, which, in turn, leads to the nuclear retention of TDP-43 and other nuclear RBPs.

This study raises important considerations for the utilization of 5EU in studies of RNA metabolism. Moreover, it confirms the dependence of TDP-43 localization on the subcellular localization of its RNA binding partners and demonstrates that increasing endogenous RNA binding sites for TDP-43 can promote its nuclear accumulation without disrupting its function.

## RESULTS

### 5EU promotes the nuclear accumulation of TDP-43 and other RBPs

TDP-43, like many predominantly nuclear RBPs, exhibits transcriptional blockade-induced translocation to the cytoplasm,^20,23,34^ which we have shown is due to depletion of nascent RNAs that tether TDP-43 in the nucleus and oppose its passive nuclear exit.^21^ To monitor changes in the subcellular localization of nascent RNAs in parallel with TDP-43, we treated HeLa cells with 5EU for 24h prior to 2 h transcriptional blockade with the selective RNA Pol II inhibitor, NVP2 (Fig. 1a). Surprisingly, cells pre-treated with 5EU showed dose-dependent nuclear accumulation of TDP-43 at steady state and striking attenuation of NVP2-induced TDP-43 nuclear exit (Fig. 1b-d). 5EU also attenuated TDP-43 nuclear exit induced by the pan-transcriptional inhibitor actinomycin D (ActD), demonstrating that the effect was not limited to NVP2 (Supplementary Fig. 1a-c). 5EU-induced TDP-43 nuclear accumulation was not observed after 1, 3, or 6h, and required a minimum of 12h 5EU pretreatment (Supplementary Fig. 1a). To exclude a batch-specific effect, we compared 5EU from three different commercial sources (≥95% pure by HPLC) and found that all attenuated ActD-induced TDP-43 nuclear exit (Supplementary Fig. 1b-c). Next, we evaluated the structural specificity by comparing 5EU to uridine alone versus 5-methyl-, 5-ethyl-, and 5-vinyl-uridine (Fig. 1e-g). Neither uridine alone nor the other 5-carbon-substituted analogs altered TDP-43 localization at steady state or after NVP2 treatment.

**Figure 1.**
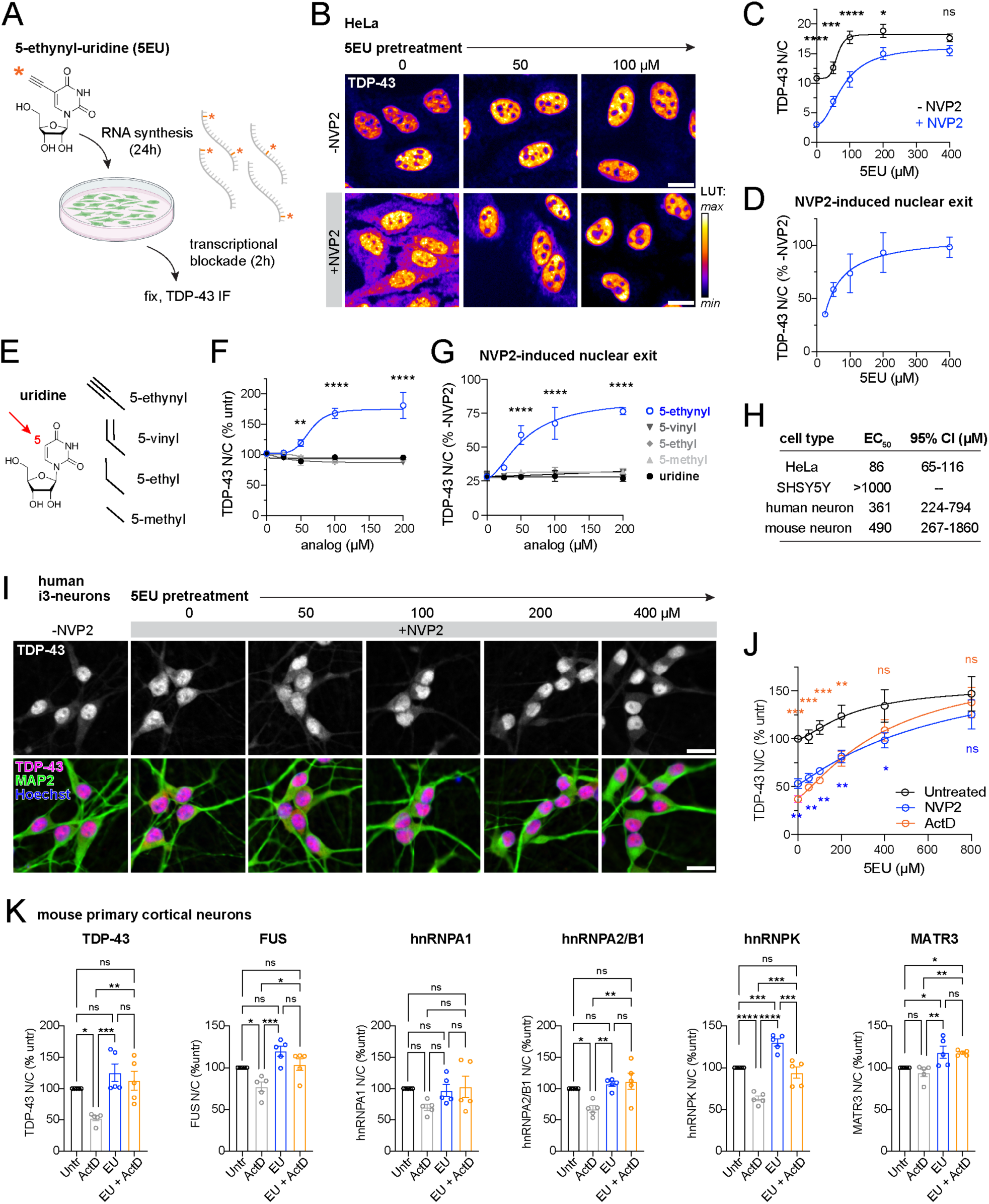
5EU promotes TDP-43 nuclear accumulation in cell lines and human neurons. A. Schematic of the nuclear retention assay, in which 5EU (*) is incorporated into newly synthesized RNAs of living cells for 24 h, followed by washout and experimental perturbation (here, 2 h transcriptional blockade with NVP2). B. Representative TDP-43 immunofluorescence images of HeLa cells pretreated with 5EU at indicated doses for 2 4h, followed by 5EU washout +/- 2 h pulse of 250 nM NVP2. The intensity histogram for each image was spread between the dimmest and brightest pixels, and a pseudo-color linear LUT was applied (see legend). Scale bar = 20 µm. C. TDP-43 N/C in 5EU-treated HeLa cells with or without NVP2 treatment. The mean ± SD of 3 independent replicates is shown, with an average of ∼2700 cells/treatment condition/replicate. D. TDP-43 N/C from (C) expressed as NVP2-treated versus untreated, representing the NVP2-induced TDP- 43 nuclear exit at each 5EU dose, normalized to the steady state rise in nuclear TDP-43. E. Structure of uridine analogs with varying substitution at the 5-carbon (arrow), tested in (F-G). F. TDP-43 N/C (% untreated cells) following uridine analog treatment for 24 h. G. TDP-43 N/C in NVP2-treated versus untreated (-NVP2) cells, following uridine analog pretreatment for 24 h, washout, and 2 h NVP2 transcriptional blockade. In F-G, the mean ± SD of 3 independent replicates is shown, with an average of ∼2100 cells/treatment condition/replicate. H. EC_50_ for 5EU attenuation of NVP2-induced TDP-43 nuclear exit in the indicated cell types. I. Representative TDP-43 immunofluorescence images of human i3-neurons pretreated with 5EU at indicated doses for 24 h, followed by 5EU washout +/- 2 h pulse of 250 nM NVP2. The intensity histogram for each image was spread between the dimmest and brightest pixels. Overlays with nuclear (Hoechst) and cytoplasmic (MAP2) counterstains are shown. Scale bar = 20 µm. J. TDP-43 N/C (% untreated cells) in 5EU-pretreated i3 neurons with or without 2 h pulse of NVP2 or ActD. The mean ± SD of 3 independent replicates is shown, with an average of ∼2400 cells/treatment condition/replicate. K. N/C ratio of TDP-43 and other nuclear RNA-binding proteins (% untreated cells) in mouse primary cortical neurons pretreated with 100 µM 5EU for 24 h, followed by a 2 h pulse of ActD. The mean ± SD of 5 independent replicates is shown, with an average of ∼700 cells/treatment condition/replicate. In C, F, G, J, K, ns= not significant, *p<0.05, **p<0.01, ***p<0.001, ****p<0.0001 by one-(K) or two-way (C,F,G,J) ANOVA with Tukey’s post-hoc test. Asterisk colors (J) correspond to treatment condition (vs. untreated cells).

Next, we compared 5EU-induced TDP-43 nuclear accumulation across multiple cell types (summarized in Fig. 1h). Human induced pluripotent stem cell (iPSC)-derived cortical neurons (i3-neurons, Fig. 1i-j) and mouse primary cortical neurons (Supplementary Fig. 1c) both demonstrated dose-dependent 5EU-induced TDP-43 nuclear accumulation, albeit with higher 24h EC_50_ than HeLa cells (490 and 361 µM vs. 66 µM). As in HeLa cells, 5EU pretreatment attenuated TDP-43 nuclear exit in human i3-neurons induced by either NVP2 or ActD (Fig. 1j). SHSY5Y human neuroblastoma cells did not display 5EU-induced TDP-43 nuclear accumulation at any dose tested (up to 1mM). ‘Click’ labeling of 5EU-treated cells after 24h showed robust labeling of HeLa cells and i3-neurons but only trace labeling in SHSY5Y cells at the highest doses, suggesting that slowed 5EU uptake or incorporation in SHSY5Y cells accounts for the lack of response in this cell line (Supplementary Fig. 2a-d). 5EU exerted marked antimitotic effects in HeLa cells, where a time-dependent decrease in cell count was observed, in parallel with a decline in metaphase cells (Supplementary Fig. 3a-c). No decrease in cell number was observed in either human or mouse cortical neurons to suggest 5EU-induced cell death (Supplementary Fig. 3d-e).

To test if the effect of 5EU was specific for TDP-43 or also affected other nuclear RBPs, we treated mouse primary neurons (Fig. 1k) and HeLa cells (Supplementary Fig. 4a-b) with 5EU for 24h, pulsed with ActD or NVP2, respectively, and fixed and immunostained cells for a panel of other RBPs. Like TDP-43, 5EU attenuated the transcriptional blockade-induced nuclear exit of multiple other RBPs, including FUS, hnRNPA2/B1, and hnRNPK in primary neurons, and FUS, hnRNPA2/B1, and ELAVL1/HuR in HeLa cells. As we previously reported,^21^ Matrin-3 does not exhibit nuclear egress after transcriptional blockade, and its localization was minimally affected by 5EU in both cell types. Finally, we probed for any 5EU-induced alteration in RBP expression by Western blot (Supplementary Fig. 4c-d) and saw no significant changes in the abundance of TDP-43 or the other RBPs tested after 24h 5EU treatment. Thus, 5EU promotes the nuclear accumulation of TDP-43 and related RBPs in a time- and dose-dependent manner, in HeLa cells, mouse primary neurons, and human iPSC-derived

### 5EU attenuates TDP-43 mislocalization due to disruption of the nuclear transport apparatus

To test if 5EU-induced TDP-43 nuclear accumulation occurs in contexts outside of transcriptional blockade, we tested two models of nucleocytoplasmic transport (NCT) disruption pertinent to motor neuron disease pathogenesis (Fig. 2). Studies in mutant *C9ORF72*-ALS/FTD^35^ and TDP-43 aggregation models^36^ suggest that disruption of RanGAP1, the Ran GTPase-activating protein, is involved in the pathogenesis of NCT defects. Recently, we showed that DLD1 cells, in which an auxin-inducible degron (AID) drives the rapid degradation of RanGAP1, display marked cytoplasmic mislocalization of TDP-43.^27^ Remarkably, pretreatment with 5EU for 12 h prior to RanGAP1 degradation conferred dose-dependent attenuation of TDP-43 mislocalization (Fig. 2a-b). Next, we generated a HeLa cell line stably expressing doxycycline-inducible charged multivesicular body protein 7 (CHMP7), which is involved in nuclear pore surveillance. The accumulation of CHMP7 in affected neurons in *C9ORF72-* and sporadic ALS has been implicated in causing NPC disruption in disease.^37^ Doxycycline-induced CHMP7-FLAG expression induced significant TDP-43 cytoplasmic mislocalization by 24h, which was attenuated by cotreatment with 5EU in a dose-dependent manner (Fig. 2c-e). Therefore, 5EU attenuated TDP-43 mislocalization induced by perturbation of the Ran gradient and structural disruption of the NPC, suggesting a generalized mechanism for promoting TDP-43 nuclear retention.

**Figure 2.**
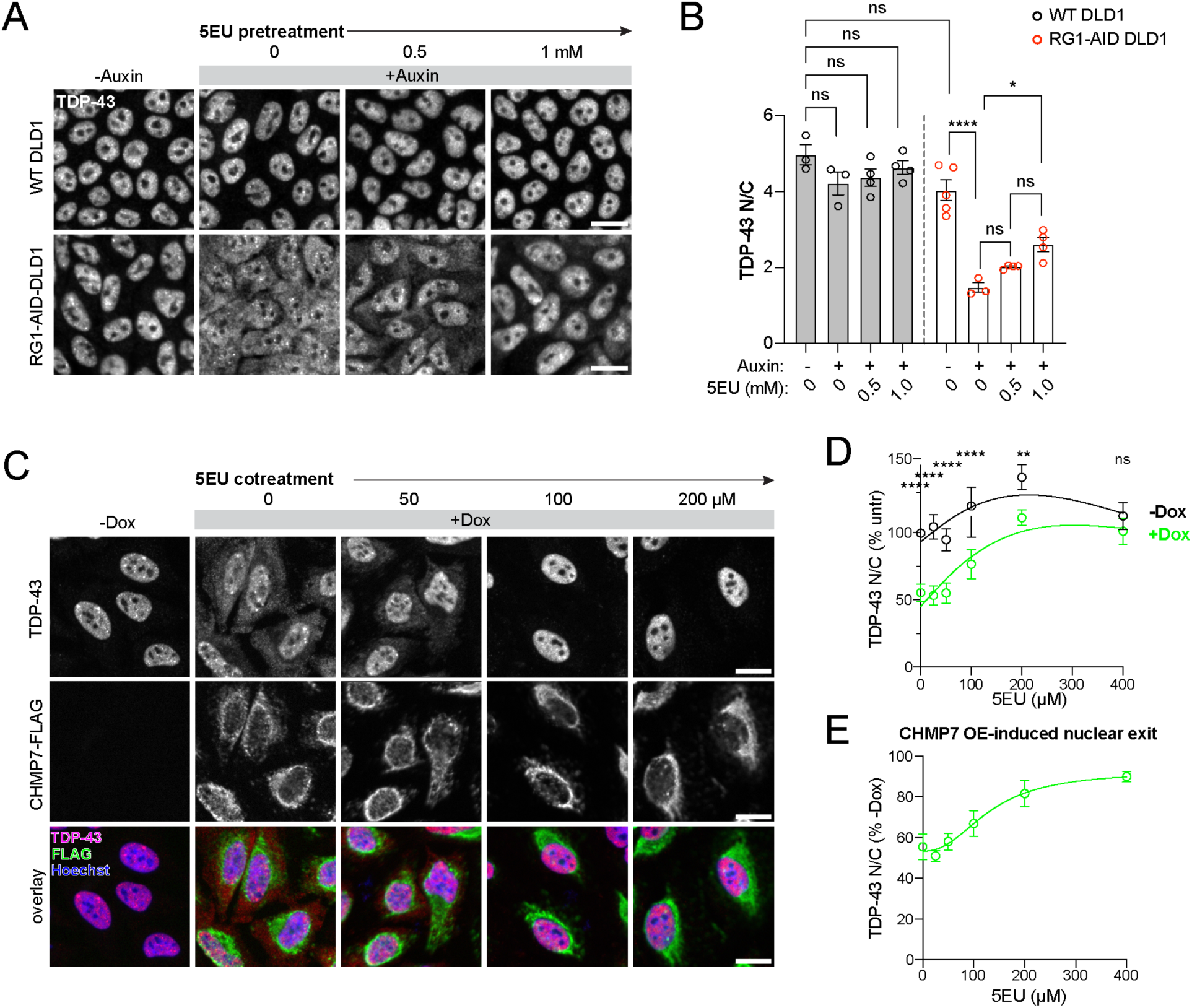
5EU attenuates TDP-43 mislocalization due to disruption of the NCT apparatus. A. Representative TDP-43 immunofluorescence staining in wild-type (WT) versus RanGAP1 (RG1)-AID DLD1 cells pretreated with the indicated dose of 5EU for 12 h before RG1 ablation by treatment with 500 µM auxin for 2 h. Scale bar = 10 µm. B. TDP-43 N/C in untreated vs. 5EU-treated cells according to cell line and auxin exposure. The mean ± SD of 3-5 independent replicates is shown, with an average of ∼700 cells/treatment condition/replicate. C. Representative TDP-43 and FLAG immunofluorescence staining in doxycycline (Dox)-inducible CHMP7-FLAG overexpressing (OE) HeLa cells. Cells were treated with Dox +/- 5EU cotreatment at the indicated doses for 24 h. Scale bar = 20 µm. D. TDP-43 N/C (% untreated cells) in +/- Dox-induced cells cotreated with 5EU. The mean ± SD of 4 independent replicates is shown, with an average of 2200 cells/treatment condition/replicate. E. TDP-43 N/C from (D) expressed as Dox-treated versus untreated, representing the CHMP7 OE-induced TDP-43 nuclear exit at each 5EU dose. In A,C, the intensity histogram for each image was spread between the dimmest and brightest pixels. In B,D, ns= not significant, *p<0.05, **p<0.01, ***p<0.001, ****p<0.0001 by one-(B) or two-way (D) ANOVA with Tukey’s post-hoc test.

### 5EU promotes RNA-binding-mediated TDP-43 nuclear retention

Since 5EU is incorporated into nascent RNAs, we hypothesized that 5EU-induced TDP-43 nuclear retention depends on changes in TDP-43-RNA interactions rather than on putative non-RNA effects of 5EU. To test this, we generated monoclonal HeLa cell lines stably expressing point mutations in the TDP-43 RNA recognition motifs (RRM) that abolish RNA binding, including 2KQ^38^ and 5FL^39^ (Fig. 3a). As we previously reported in transiently transfected cells,^21^ V5-tagged TDP43-2KQ and TDP43-transcriptional blockade (Fig. 3b-c). Similarly, 24 h treatment with 100 µM 5EU did not induce any change in TDP43-2KQ and TDP43-5FL localization, in contrast to wild-type TDP-43. Thus, TDP-43 nuclear sequestration induced by 5EU requires TDP-43 RRM-mediated binding to RNA.

**Figure 3.**
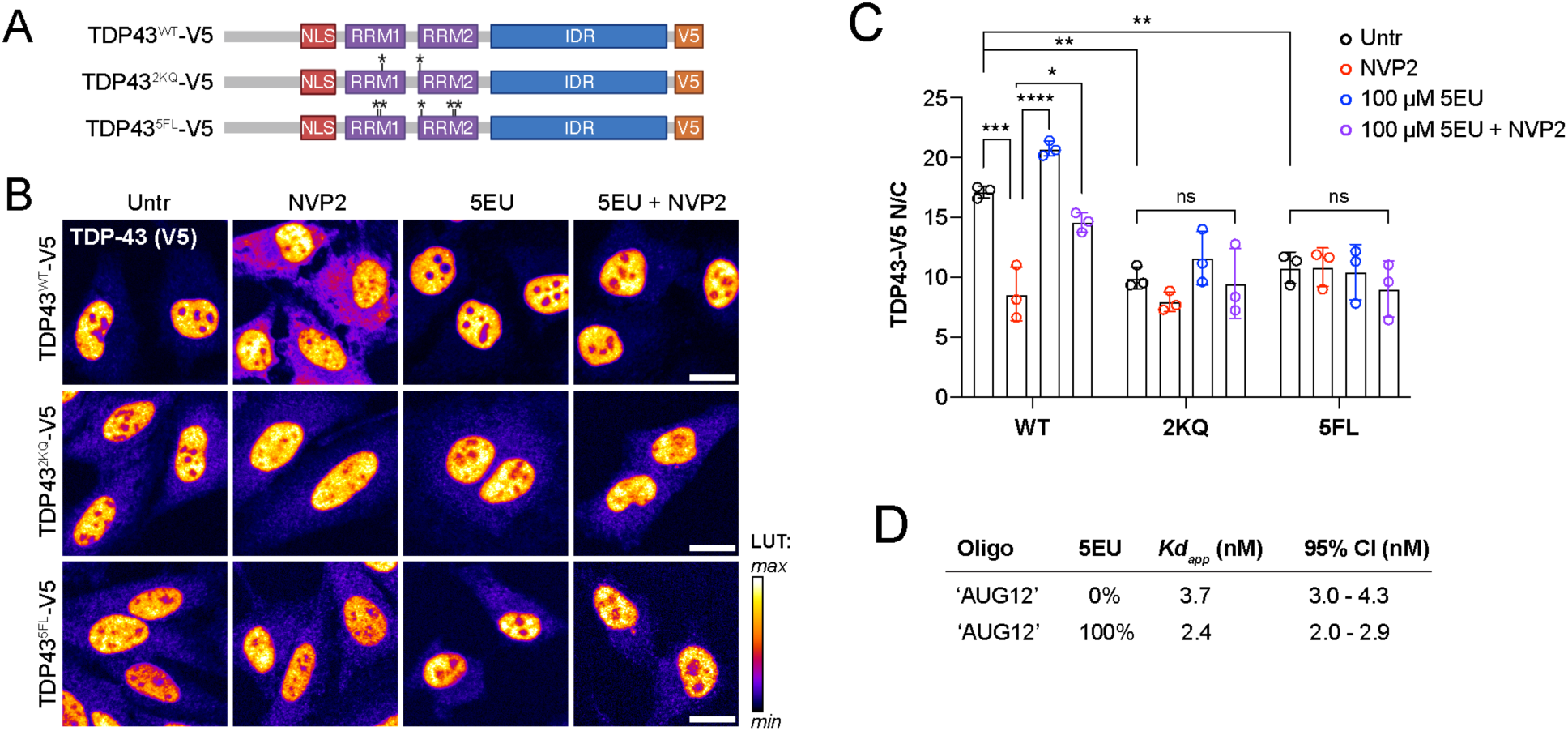
5EU promotes RNA binding-mediated TDP-43 nuclear accumulation. A. V5-tagged TDP-43 RNA recognition motif (RRM) point mutation (*) constructs. B. Representative V5 immunofluorescence images from HeLa cells stably expressing the constructs in (A), pretreated with 100 µM 5EU for 24 h, followed by 5EU washout +/- 2 h pulse of 250 nM NVP2. The intensity histogram for each image was spread between the dimmest and brightest pixels and a pseudo-color linear LUT was applied (see legend). Scale bar = 20 µm. C. TDP43-V5 N/C ratio according to cell line and treatment. The mean ± SD of 3 independent replicates is shown, with an average of ∼2300 cells/treatment condition/replicate. NS= not significant, *p<0.05, **p<0.01, ***p<0.001, ****p<0.0001 by two-way ANOVA with Tukey’s post-hoc test. D. TDP-43 binding affinity (*Kd_app_*) for *in vitro-*transcribed oligonucleotides containing 0% or 100% 5-ethynyl-modified uridine.

Next, we hypothesized that 5EU incorporation into endogenous RNAs could augment TDP-43 nuclear localization by at least two alternative mechanisms: 1) direct effects of 5EU on TDP-43-RNA binding, or 2) indirect effects of 5EU on the nuclear/cytoplasmic localization of TDP-43 RNA targets. To test for direct effects on TDP-43-RNA binding, we analyzed TDP-43 binding affinity for unmodified versus 5EU-modified synthetic RNA oligonucleotides. We *in vitro* transcribed ‘AUG12’ (5’-GUGUGAAUGAAU-3’), a motif that stably binds the TDP-43 RRM1-2 domains,^40^ with either 0% or 100% 5EU and analyzed TDP-43 binding affinity by measuring changes in TDP-43 intrinsic fluorescence upon RNA binding (Fig. 3d, Supplementary Fig. 5a-b).^41^ TDP-43 affinity for unmodified ‘AUG12’ was 3.7 nM (95% CI 3-4.3 nM).100% 5EU incorporation, far exceeding the estimated 5.7% 5EU incorporation into endogenous mRNAs,^32^ only subtly increased the TDP-43 RNA binding affinity (*Kd_app_* 2.4 nM, 95% CI 2-2.9 nM). Thus, altered TDP-43-RNA binding is unlikely to explain the robust 5EU-induced TDP-43 nuclear retention in living cells.

### 5EU induces nuclear RNA accumulation

We previously demonstrated that TDP-43 nuclear localization is coupled to nuclear GU-rich pre-mRNA binding and abundance.^21,27^ Since 5EU-induced TDP-43 nuclear sequestration requires RRM-mediated RNA binding (Fig 3a-c), but extreme (supraphysiological) 100% 5EU substitution causes only subtle changes in TDP-43-RNA binding affinity (Fig. 3d), next we investigated indirect effects of 5EU on the localization and abundance of RNA. To evaluate 5EU-induced changes in nuclear polyadenylated RNAs, HeLa cells were treated with increasing doses of 5EU for 24 h, with or without a subsequent 2 h pulse of NVP2, and then fixed for polyA-FISH (Fig. 4a-d). We observed a striking, dose-dependent increase in the polyA-N/C ratio (Fig. 4b), driven by a significant increase in nuclear indicating a possible 5EU-induced blockade in mRNA export. 2 h transcriptional blockade with NVP2 did not significantly alter the polyA signal (Fig. 4b). Next, we repeated the RNA-FISH with a (CA)6-Cy5 probe to label GU-repeat motifs within TDP-43 RNA binding sites (Fig. 4e-g). Interestingly, the resulting (GU)6-FISH staining pattern was almost exclusively nuclear and exhibited a nucleolar sparing pattern reminiscent of TDP-43 localization (Fig. 4e). Unlike the polyA-FISH, the (GU)6-FISH signal markedly decreased after 2 h NVP2 treatment, consistent with the expected location of GU-repeat clusters within short-lived intronic RNA segments of nascent pre-mRNAs. Remarkably, 5EU pretreatment attenuated the NVP2-induced loss of the (GU)6-FISH signal in a dose-dependent manner (Fig.4g), suggesting that 5EU preferentially stabilizes the nuclear pool of GU-rich pre-mRNAs. To verify the putative stabilization of GU-rich RNAs by 5EU, HeLa cells were treated for 24h with 100 µM 5EU, followed by washout and (GU)6-FISH labeling after 0, 1, 2, 4, or 8 h NVP2 (Fig. 4h-j). Indeed, 5EU pretreatment significantly slowed the transcriptional blockade-induced loss of (GU)6-RNAs, approximately doubling the half-life of the nuclear (GU)6-FISH signal (Fig. 4j).

**Figure 4.**
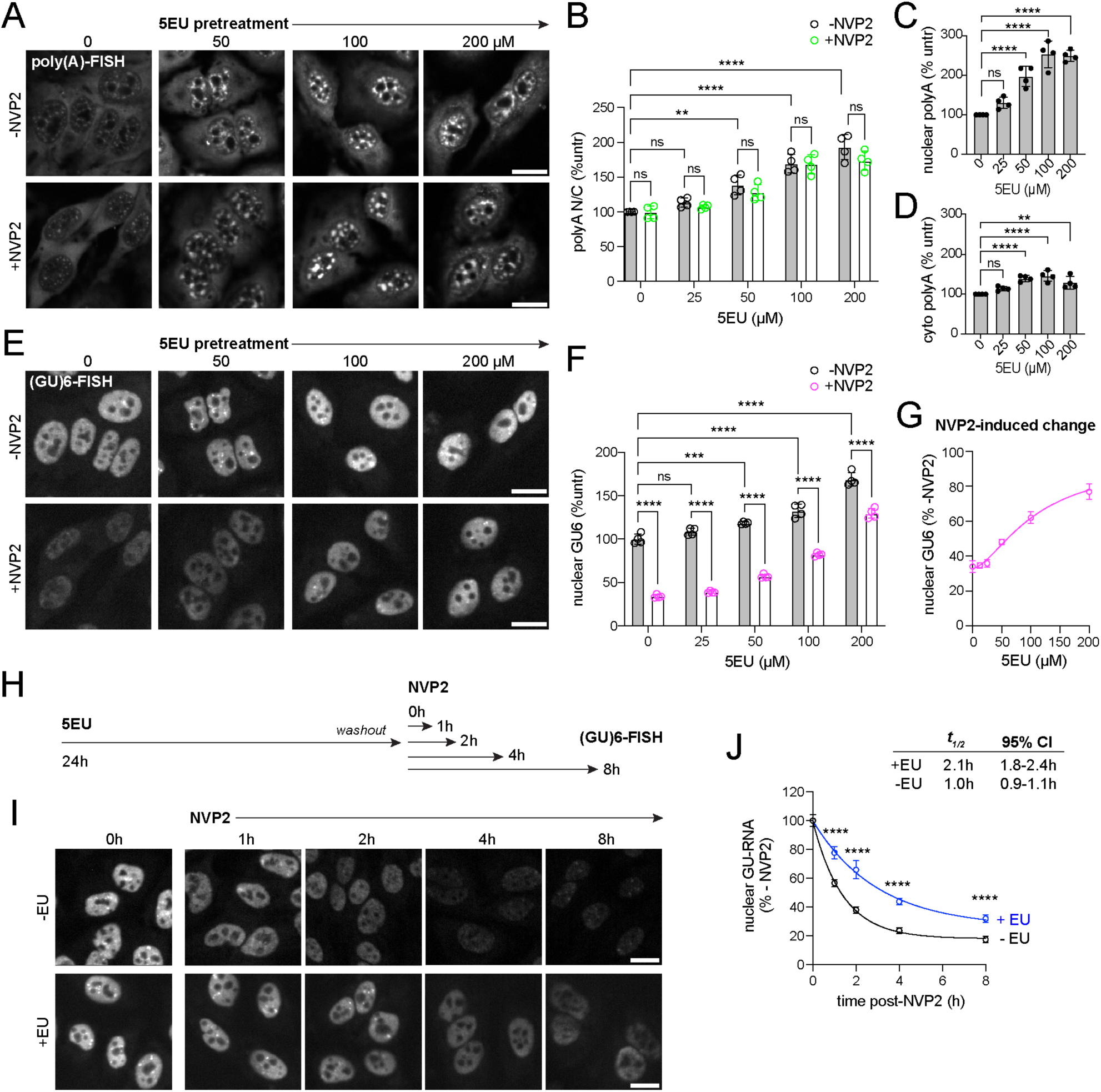
5EU induces the accumulation of nuclear RNAs. A. Representative poly(A)-FISH (probe = oligo(dT)-FAM) images from HeLa cells pretreated with 5EU at indicated doses for 24 h, followed by 5EU washout +/- 2 h pulse of 250 nM NVP2. The intensity histogram for each image was normalized to the dimmest and brightest pixels of the maximally treated (200 µM 5EU) cells. Scale bar = 20 µm. B. Poly(A) N/C ratio expressed as % untreated cells. The mean ± SD of 4 independent replicates is shown, from an average of ∼2500 cells/treatment condition/replicate. C-D. Nuclear (C) and cytoplasmic (D) poly(A) intensity corresponding to the data in (B), expressed as % untreated cells. E. Representative (GU)6-FISH (probe = CA6-Cy5) images from HeLa cells pretreated with 5EU for 24 h, followed by 5EU washout +/- 2 h pulse of 250 nM NVP2. The intensity histogram for each image was normalized to the dimmest and brightest pixels of the maximally treated (200 µM 5EU) cells. Scale bar = 20 µm. F. Nuclear (GU)6 intensity expressed as % untreated cells. The mean ± SD of 4 biological replicates is shown, from an average of ∼2500 cells/treatment condition/replicate. G. Nuclear (GU)6 intensity from (F) in NVP2-treated vs untreated (-NVP2) cells, representing the NVP2-induced change in nuclear (GU)6-containing RNA motifs. H. Schematic of nuclear (GU)6-RNA-decay experiment in which cells were pretreated with 100 µM 5EU for 24 h prior to washout and NVP2 transcriptional blockade for 0, 1, 2, 4, or 8 h prior to (GU)6-RNA FISH. I. Representative (GU)6-FISH images from the experiment diagrammed in H. The intensity histogram for each image was normalized to the dimmest and brightest pixels of the corresponding - EU or +EU-treated cells at time 0. Scale bar = 20 µm. J. Nuclear (GU)6 intensity in NVP2-treated vs untreated (-NVP2) cells, representing the NVP2-induced change in nuclear (GU)6-containing RNA motifs at each timepoint. The half-life (*t_1/2_*) of the nuclear (GU)6 signal for 5EU-treated vs. untreated cells is shown. The mean ± SD of 5 biological replicates is shown, from an average of ∼1200 cells/treatment condition/replicate. In B, F, J, ns= not significant, **p<0.01, ****p<0.0001 by two-way ANOVA with Tukey post-hoc test.

Given the >12 h delay in 5EU - induced effect on TDP43 nuclear accumulation (Supplementary Fig. 1a), we monitored how the TDP-43 localization compares with the nuclear incorporation of the 5EU-modified RNAs. ‘Click’ labeling in 5EU-treated HeLa cells after 0, 1, 6, and 24h of treatment showed a time-dependent increase in nuclear 5EU-RNA that approximately doubled from 1 to 6 h, and doubled again by 24 h, compared to a slower increase in the cytoplasmic signal (Supplementary Fig. 6a-b). Nuclear export of 5EU-RNA was detected by the appearance of a cytoplasmic signal and a progressive decrease in the N/C ratio over time. PolyA-FISH and (GU)6-FISH also showed time-dependent nuclear accumulation (Supplementary Fig. 6c-d). Together, these findings demonstrate that 5EU induces a time- and dose-dependent increase in nuclear RNAs. Moreover, the 5EU-induced attenuation of transcriptional blockade-induced TDP-43 nuclear exit is paralleled by apparent stabilization of nuclear GU-RNAs.

### 5EU preserves nuclear TDP-43 RNA binding and function

To further investigate the effect of 5EU on TDP-43 RNA binding, HeLa cells were treated for 24 h with 100 µM 5EU, with or without a 2 h pulse of NVP2, and then UV crosslinked and harvested for TDP-43 eCLIP.^42^ Consistent with previous reports,^2,3^ in untreated cells the TDP-43 peaks were predominantly found in introns, particularly within distal introns (Fig. 5a). 2 h transcriptional blockade with NVP2 caused a marked decrease in intronic binding sites and a concomitant increase in peaks found in mRNA 3’-UTR and coding sequences, suggesting preferential TDP-43 binding to cytoplasmic mRNAs following NVP2 treatment (Fig. 5a). 5EU treatment alone did not markedly change the distribution of TDP-43 RNA binding sites. However, in cells pre-treated with 5EU for 24 h, followed by 2 h NVP2, 5EU abolished the NVP2-induced shift from intronic to mRNA binding sites (Fig. 5a). To explore individual peaks, we considered whether peaks observed in untreated cells were altered upon either NVP2 or 5EU treatment. As above, peak-level TDP-43 RNA binding in 5EU pre-treated cells exposed to NVP2 was nearly identical to NVP2-untreated cells, whereas a dramatic loss in signal was seen in NVP2-treated cells without 5EU pre-treatment (Fig. 5b, Supplementary Fig 7a). 5EU pretreatment preserved TDP-43 binding to distal introns within hundreds of unique transcripts, suggesting a transcriptome-wide effect rather than an effect mediated by a narrow subset of transcripts. Selected examples of 5EU-induced preservation of TDP-43 binding to intronic, GU-rich regions are shown (Fig. 5c). In all conditions, unbiased motif analysis by HOMER indicated the top TDP-43 binding motifs were the expected GU- and GUA-rich sequences (Fig 5d). K-mer analysis showed highly correlated enrichment across all conditions (Fig. 5e-g), with both GU-rich and GAAUG motifs enriched similarly across all treatments (Supplementary Fig 7b-c).

**Figure 5.**
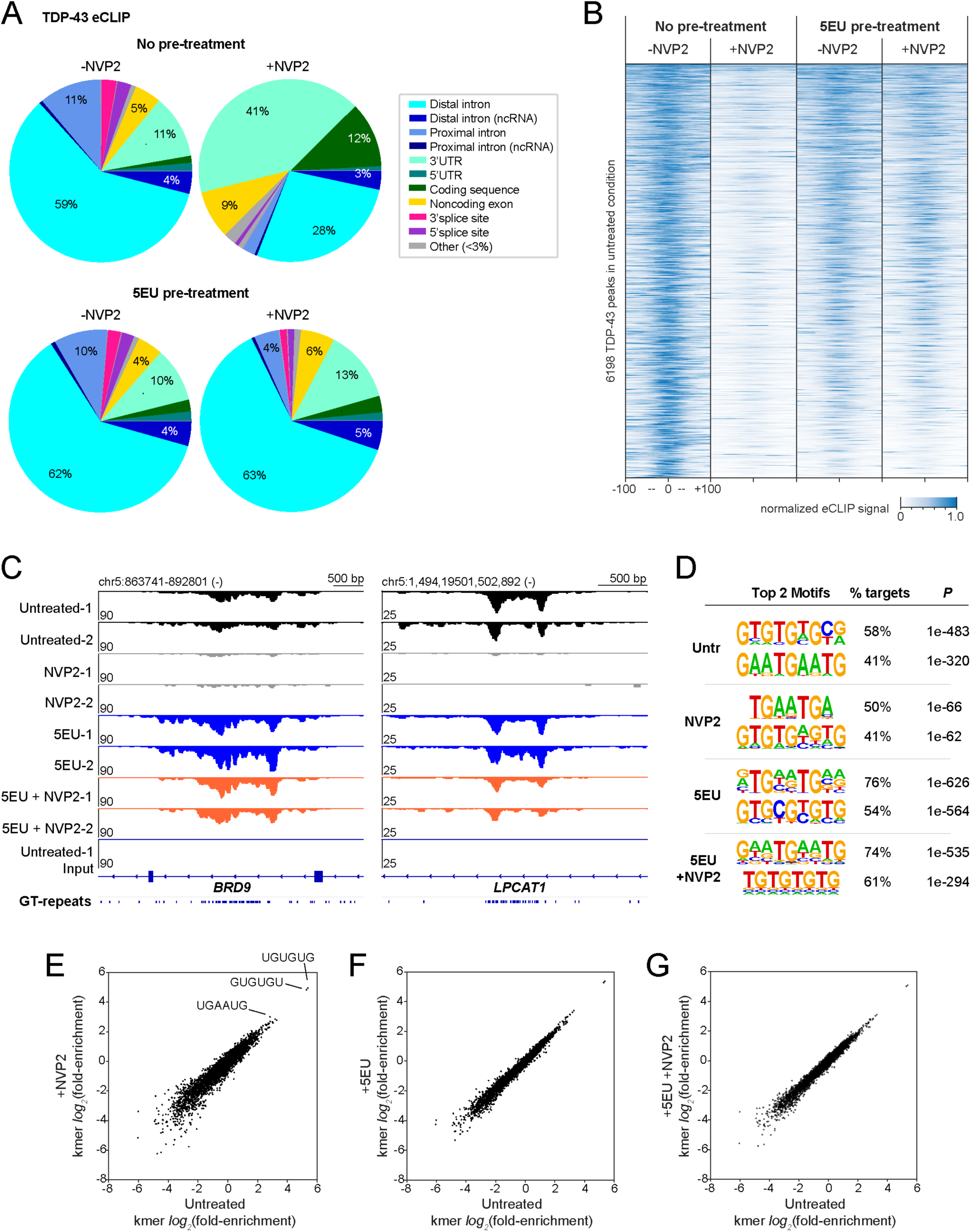
5EU prevents transcriptional blockade-induced depletion of TDP-43 intronic binding sites. A. Distribution of mapped peaks from TDP-43 eCLIP in untreated HeLa cells versus cells pretreated with 100 µM 5EU for 24 h, followed by 5EU washout ± 2 h pulse of 250 nM NVP2. N=2 independent replicates. B. 200nt windows centered on each TDP-43 eCLIP peak in untreated vs. 5EU ± NVP2-treated cells. Each row indicates the normalized read density. Replicate 1 (shown here) and Replicate 2 (Supplementary Fig. 7A) were normalized separately to control for batch effects. C. Representative maps of TDP-43 binding within the distal intron of two transcripts, *BRD9* and *LPCAT1,* across treatment conditions and replicates. The input from untreated cell sample 1 is provided for comparison, and GT-repeat clusters within the primary sequence are shown. D. The top 2 TDP-43 binding motifs identified by HOMER across treatment conditions. E-G. Log_2_(fold-enrichment) for individual k-mers observed in significant, reproducible peaks for TDP-43 eCLIP performed in (x-axis) untreated cells versus cells treated with (y-axis) NVP2 (E), 5EU (F), or 5EU followed by NVP2 (G).

TDP-43 is a prolific splicing regulator that, among other functions, critically represses cryptic exons.^4–6^ Because 5EU did not alter the distribution of TDP-43 binding sites within introns, we predicted that 5EU would not alter TDP-43 splicing function. To investigate, total RNA sequencing was performed in untreated vs. 100 µM 5EU-treated cells (24 h). The data were compared to published data from TDP-43 siRNA-treated HeLa cells,^4^ to probe for any failure of TDP-43 cryptic single locus, *FAM114A2*, suggesting that 5EU does not broadly disrupt TDP-43 function (Fig. 6a). To further validate this, we performed RT-PCR analysis of three cryptic exon splicing events that are sensitive to partial loss of TDP-43 function, *EPB41L4A, ARHGAP32,* and *POLDIP3* (Fig. 6b). None showed 5EU-induced failure of cryptic exon repression. Thus, 5EU preserves TDP-43 nuclear RNA binding and splicing regulatory function.

**Figure 6.**
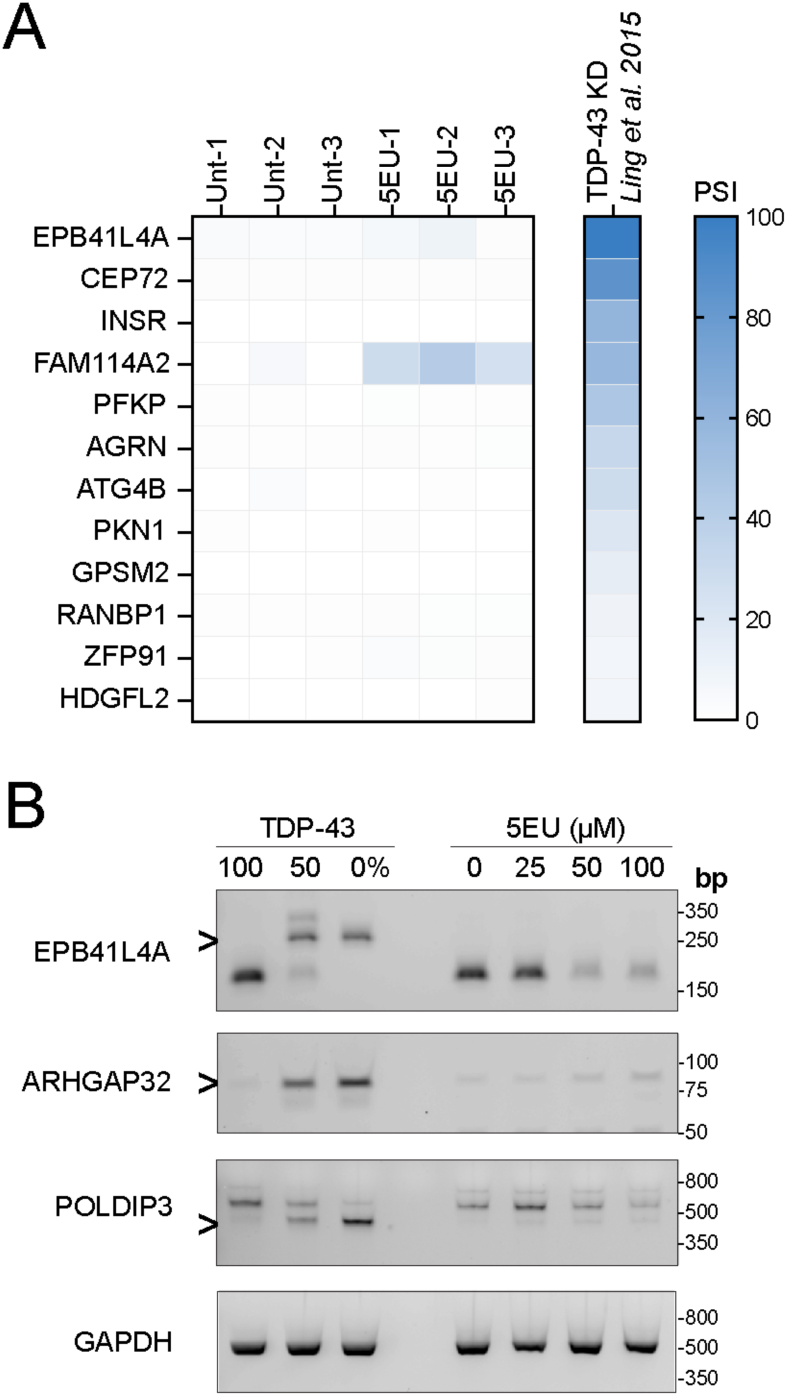
TDP-43 repression of cryptic exons is preserved in 5EU-treated cells. A. PSI of TDP-43-regulated cryptic exons in the indicated transcripts, compared to published data set from TDP-43 siRNA-treated HeLa cells.^4^ N=3 independent replicates. B. RT-PCR of TDP-43-regulated cryptic exons (>) in *EPB41L4A, ARHGAP32, and POLDIP3,* in HeLa cells treated with the indicated doses of 5EU for 24h. Controls for TDP-43 expression level include untreated cells (100%), TDP-43 siRNA-treated cells, 24 h (50%), and TDP-43 CRISPR knockout cells (0%).

### 5EU does not markedly alter RNA decay

Based on the data from GU-RNA FISH (Fig. 4) and TDP-43 eCLIP (Fig. 5) that 5EU attenuates the transcriptional blockade-induced loss of GU-rich nuclear RNAs and preserves TDP-43 binding to intronic sites, we hypothesized that 5EU incorporation may prolong RNA half-life. To test this, we performed an RNA decay analysis, in which HeLa cells were treated with 100 µM 5EU for 24 h, followed by washout and transcriptional blockade with NVP2 for 0, 1, 2, 4, or 8 h prior to RNA extraction and total RNAseq (Fig. 7a). Transcript abundance at each timepoint was normalized to spike-in controls, and further normalized to time 0, to construct RNA decay curves and determine transcript half-life (Fig. 7b). Population-wide analysis suggested a modest 5EU-induced prolongation of RNA half-life (median ∼9 h for untreated cells, 10 h for 5EU-treated cells), with a wider distribution of half-lives in the untreated samples, but unsupervised clustering did not identify any major changes in half-life within specific RNA subsets (Fig. 7c-e). Thus, our findings are unlikely to be explained by 5EU-induced prolongation of RNA half-life.

**Figure 7.**
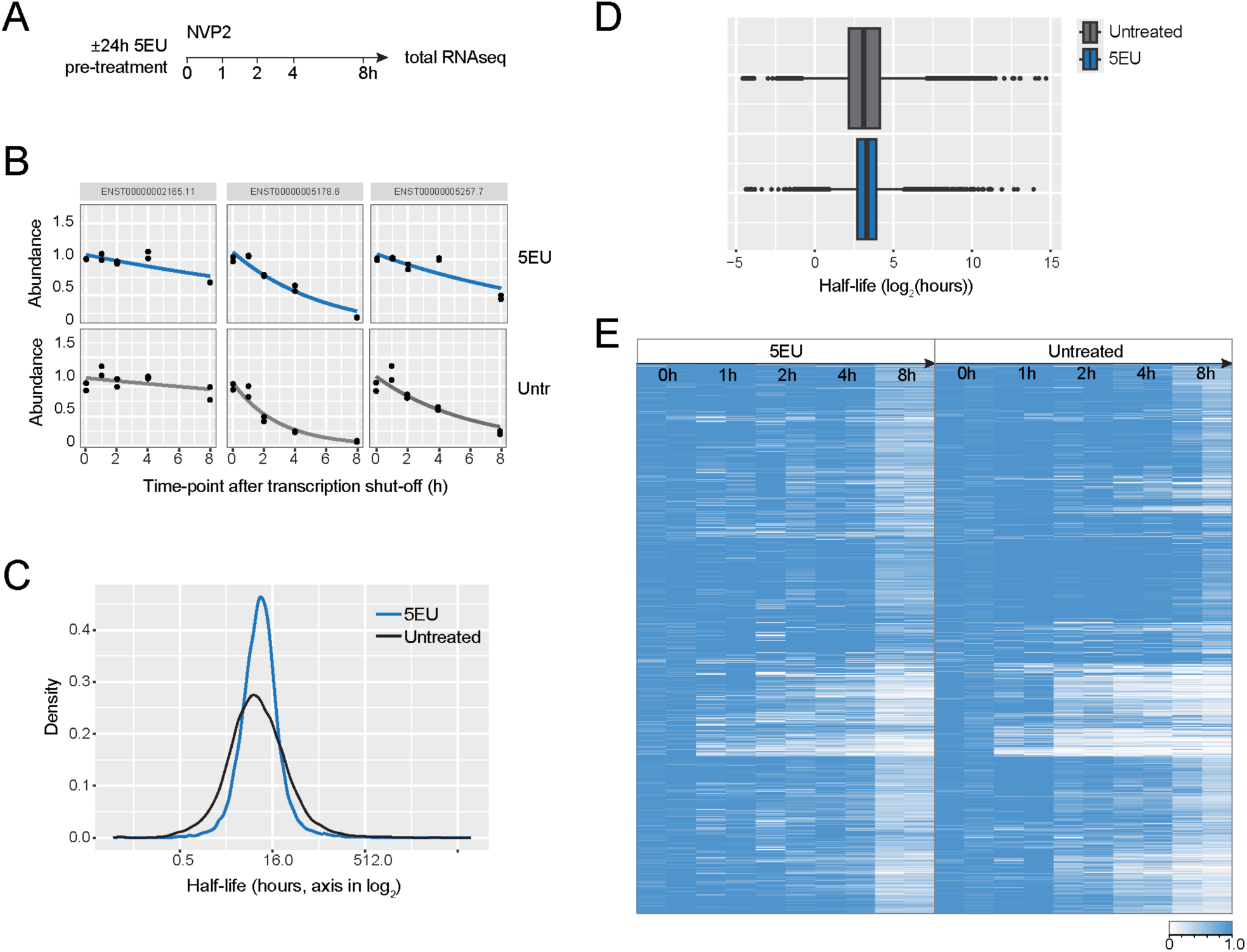
5EU does not markedly prolong RNA half-life. A. Schematic of RNA decay assay, in which a subset of HeLa cells was treated with 100 µM 5EU for 24 h, followed by washout and transcriptional blockade with NVP2 for 0, 1, 2, 4, or 8 h prior to RNA extraction and total RNAseq. B. Sample RNA decay curves for individual transcripts. C. Distribution of RNA half-lives (log_2_) in 5EU-treated vs untreated cells. D. Boxplot representing the distribution of RNA half-lives (log_2_) in 5EU-treated vs untreated cells. E. Heatmap of transcript abundance (TPM) at each timepoint in 5EU-treated vs. untreated cells, normalized to time 0. In B-E, n=2 technical replicates. The abundance of all transcripts was first normalized to RNA spike-in controls.

### 5EU alters gene expression and alternative splicing

To further investigate changes in nuclear RNA processing that might contribute to 5EU-induced nuclear polyA and GU-rich RNA accumulation, we performed differential gene expression and alternative splicing analysis in untreated HeLa cells versus cells treated with 100 µM 5EU for 24 h (Fig. 8). Principal component analysis showed distinct clustering of untreated versus 5EU-treated samples (Fig. 8a). Differential gene expression analysis showed numerous significantly up- and down-regulated transcripts (Fig. 8b). Consistent with our immunoblots (Supplementary Fig. 4c-d), TDP-43 (*TARDBP*) expression was unchanged at the transcript level. Gene ontology (GO) analysis of up- and down-regulated genes showed prominent upregulation of genes in RNA processing, splicing, and ribonucleoprotein (RNP) complex biogenesis (Fig. 8c). Meanwhile genes involved in functions such as cytoskeletal organization and cell motility were downregulated. Consistent with alterations in nuclear RNA metabolism, immunostaining for Cajal bodies, nuclear speckles, and nucleoli in 5EU-treated cells showed dose-dependent changes in number and size (Supplementary Fig. 8a-f).

**Figure 8.**
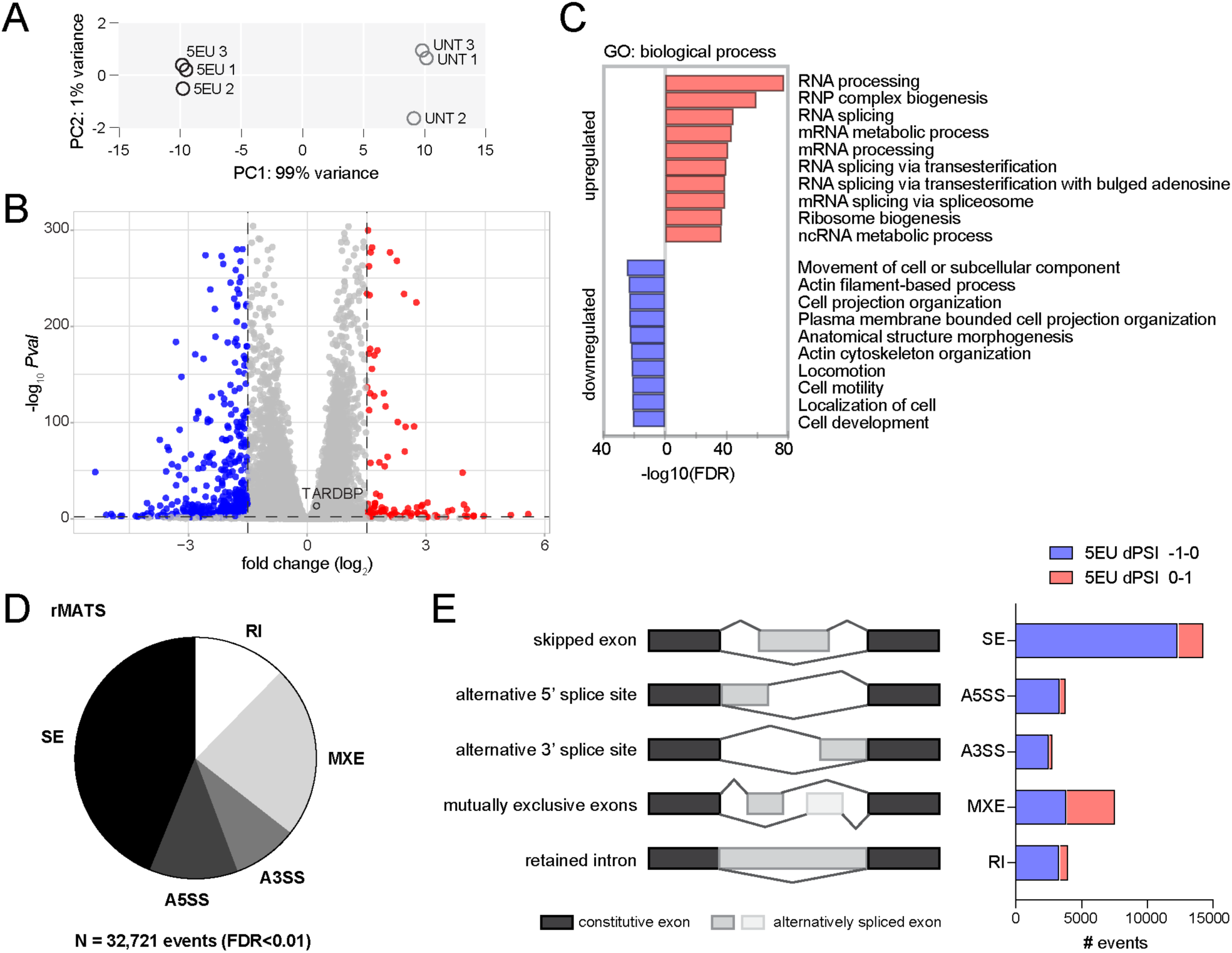
5EU alters gene expression and RNA splicing. A. PCA plot of 5EU-treated (100 µM, 24 h) versus untreated HeLa cells from total RNAseq. N=3 independent replicates/group. B. Volcano plot of up- and down-regulated genes in 5EU-treated cells. Of note, there was no significant change in the expression of *TARDBP* as indicated. C. Gene ontology (GO) analysis of the top 10 up- and down-regulated biological processes in 5EU-treated cells, as ranked by the false discovery rate (FDR). D. Alternative splicing events in 5EU-treated versus untreated cells from rMATS analysis, according to type of event (RI = retained intron, SE = skipped exon, MXE = mutually exclusive exons, A5SS = alternative 5’ splice site, A3SS = alternative 3’ splice site). E. Categorization of events in (D) according to dPSI > or < 0 by treatment category.

Next, replicate multivariate analysis of transcript splicing (rMATS)^43^ was performed to analyze alternative splicing (Fig. 8d-e). rMATS detected >30,000 differential splicing events in 5EU-treated versus untreated cells (FDR≤0.01), primarily at skipped exon (SE) and mutually exclusive exons (MXE), but also a subset of differential retained intron (RI) and alternative 5’ splice site (A5SS) or alternative 3’ splice site events (A3SS). These data suggest a significant effect of 5EU on pre-mRNA splicing, potentially affecting multiple splicing events in RNA molecules in parallel. Across all categories, the 5EU delta PSI (dPSI=PSI_5EU_-PSI_UNT_) for most events was <0 (Fig. 8e). For MXE, A5SS, and A3SS, the algorithm measures the ratio between two splice site choices; thus, the dPSI value does not inform about overall 5EU-induced upregulation or downregulation of splicing. However, for SE and RI, if we view the event as a 0-to-1 dial based on the inclusion level (PSI) of the alternative splicing feature (exon and intron, respectively), the 5EU dPSI <0 indicates a reduction in the intensity of the dial, which could suggest that 5EU is downregulating the underlying mechanisms enabling these processes.

To further investigate how 5EU affects splicing, next we used MntJULiP.^44,45^ MntJULiP calculates a per-sample measure of splicing complexity based on the relative contribution of the primary ‘isoform’ in a group of competing alternative introns (see Methods). On average, the splicing complexity of the untreated samples was significantly higher than the 5EU-treated samples (0.545 versus 0.508, *p*<0.01 by 2-tailed t-test), indicating a 5EU-induced reduction in splicing diversity. Based on this finding, we conclude that the splicing of alternative isoforms was comparatively repressed in 5EU-treated samples.

## DISCUSSION

In this study, we report previously unappreciated and significant consequences of cell treatment with the uridine analog 5EU. 5EU induced the accumulation of nuclear RNAs, RNA-bound TDP-43, and other RBPs. TDP-43 eCLIP showed that 5EU did not alter TDP-43 binding at predominantly intronic sites; however, RNAseq analyses revealed significant 5EU-induced changes in alternative splicing and an overall reduction in splicing diversity. These data suggest that 5EU may impede RNA splicing efficiency and subsequent nuclear RNA processing and export. The resulting 5EU-induced accumulation of endogenous nuclear RNAs, in a striking parallel with other recent experimental paradigms (discussed below), promotes the nuclear localization of TDP-43 and other RBPs with RNA-dependent nuclear/cytoplasmic localization.

### Further demonstration of RNA-dependent RBP localization

Transcriptional blockade induces the nuclear efflux of a large subset of RBPs with diverse RNA binding motifs,^21,34^ suggesting that the loss of nuclear RBP binding sites within nascent RNAs broadly disrupts nuclear RBP localization. Here, we report the spatial inverse, that 5EU-induced nuclear RNA accumulation promotes the nuclear accumulation of TDP-43 and other RBPs (Fig. 1 and Supplementary Fig. 4). The 5EU-responsive RBPs recognize a variety of RNA motifs, suggesting that 5EU broadly affects many RNA species targeting multiple RBPs.

The 5EU-induced nuclear accumulation of RNA (Fig. 4) parallels our recent finding that two small molecule inhibitors of splicing — isoginkgetin and pladienolide B — promoted the nuclear accumulation of intronic RNAs together with TDP-43 and hnRNPA1.^21^ Acute mRNA export blockade also caused nuclear polyA-RNA and TDP-43 accumulation.^21^ Using nucleocytoplasmic transport promotes TDP-43 nuclear retention by restricting its passive diffusion through NPC channels,^21^ an effect that can be augmented by the introduction of synthetic RNAs.^27^ The extent to which size-restricted diffusion within RNP complexes more broadly accounts for the RNA-dependent localization of RBPs beyond TDP-43 remains to be tested. However, 5EU-induced TDP-43 nuclear retention further demonstrates that the localization of TDP-43 is coupled to the subcellular gradient of its target RNAs.

### Cellular and molecular changes in 5EU-treated cells

Since the initial 2008 report,^32^ 5EU has been widely used to analyze RNA transcription and turnover,^46–53^ splicing,^54,55^ RNA export,^56,57^ and RNA subcellular localization,^58,59^ primarily in *in vitro* systems but also *in vivo,* including in yeast,^60^ *Xenopus* embryos,^61^ *Drosophila,*^62^ and mice.^32,52,63,64^ For *in vitro* RNA labeling, including in neurons,^46^ 5EU concentrations commonly range from 0.1-1 mM, with time courses from <1 h to 72 h. Notably, studies utilizing ‘EU-seq’, in which cells are pulsed with 5EU prior to EU-RNA click labeling, purification, and sequencing, often use short (≤1 h) 5EU pulses to isolate the nascent RNA pool.^65^ Thus the 5EU concentrations used in the current study are within the typical range, but the time required for effects on TDP-43 (12-24 h, Supplementary Fig. 1a) is longer than protocols most frequently used to analyze nascent RNA.

Reports of 5EU-induced cellular and molecular changes are limited, though few studies have compared 5EU-treated versus untreated cells. Consistent with our observation of cell cycle inhibition in HeLa cells (Supplementary Fig. 3), 5EU was reported to inhibit HEK293T cell proliferation without affecting cell viability when treated with up to 1 mM over a period of 5-72 h.^51^ A 5EU-induced decrease in cell counts was also reported in A549 cells treated with up to 500 µM 5EU for 48 h.^48^ 4-thiouridine (4sU), another widely-used RNA metabolic label, has similarly been reported to inhibit cell proliferation and caused time- and dose-dependent inhibition of rRNA synthesis and nucleolar stress.^66^ Notably, 4sU caused subtle alterations of pre-mRNA splicing in *in vitro* extracts and cell that 4sU concentration and time course should be minimized to avoid reducing splicing efficiency. No studies were identified to date reporting differential gene expression or splicing analysis in 5EU-treated versus untreated cells.

Nuclear RNA transcription, splicing, export, and decay are highly interdependent processes, and 5EU-induced perturbation at any stage could conceivably initiate a cascade of events leading to nuclear RNA and RBP accumulation. Though 5EU is taken up and incorporated into nascent RNAs in less than 1h (Supplementary Fig. 6a-b), 5EU-induced nuclear TDP-43 accumulation required at least 12 h (Supplementary Fig. 1), paralleling the rise in nuclear GU-RNAs (Supplementary Fig. 6d). Thus, 5EU-induced nuclear RNA and TDP-43 accumulation is unlikely to be an immediate result of 5EU incorporation into nascent RNA, but rather the downstream effect of a yet unknown cascade of events. RNA-FISH showed a 5EU-induced increase in both stable, polyadenylated (Fig. 4b) and short-lived, GU-rich RNAs (Fig. 4f), suggesting both nascent and processed RNAs accumulate in the nucleus of 5EU-treated cells. 5EU-labeled RNA was exported to the cytoplasm over time, ruling out a severe RNA export blockade (Supplementary Fig. 6a-b). However, further analysis is needed to probe for a more subtle delay in RNA export. Similarly, no gross delay in RNA decay was observed (Fig. 7), though testing shorter timepoints < 1 h could clarify the potential stabilization of short-lived RNAs. Strand-specific RNA seq^68^ or long-read sequencing^69,70^ could further characterize the accumulated nuclear RNAs to clarify how 5EU may affect pre-mRNA processing. Though there are no reports on the effects of 5EU on RNA structure, pre-mRNA structure has been shown to influence splicing outcomes.^71,72^ Interestingly, the abundant endogenous uridine isomer, pseudouridine, alters RNA duplex stability,^73^ and a highly-conserved pseudouridine in the U2 snRNA critically sculpts U2 snRNA binding to the intronic pre-mRNA branch site adenosine during spliceosome assembly.^74,75^ Given the 5EU-induced changes in alternative splicing (Fig. 8) and overall reduced splicing diversity in 5EU-treated cells, further investigation is warranted to test the hypothesis that 5EU incorporation may similarly alter pre-mRNA secondary structure or spliceosome dynamics. Pending further mechanistic clarification, our findings suggest that short pulses of low-dose 5EU should be utilized for studies investigating nuclear RNA metabolism to avoid confounding effects.

### Insights for TDP-43 modulation

The key role of RNA in regulating TDP-43 localization and solubility suggests that the disruption of RNA metabolism could contribute to TDP-43 pathology in neurodegeneration and that RNA-based interventions may promote TDP-43 homeostasis. Indeed, we recently found that transient transfection of multivalent GU-rich oligonucleotides to increase nuclear RNA binding sites for TDP-43 promoted TDP-43 nuclear localization in cultured cells.^27^ GU-rich oligonucleotides also attenuated TDP-43 aggregation in human neurons.^76^ Though oligonucleotide-based approaches enable selective, RNA motif-based targeting of TDP-43, we observed that pure GU-repeat oligonucleotides synthesized with protective phosphorothioate (PS)-bonds caused unwanted inhibition of TDP-43 function, likely by high affinity binding and displacement of TDP-43 from endogenous RNAs.^27^ TDP-43 loss of function was markedly reduced using a lower-affinity interspersed sequence (Clip34nt) based on the endogenous TDP-43 binding site in its own 3’-UTR. In the current study, the 5EU-induced accumulation of diverse, endogenous RNAs promoted TDP-43 nuclear accumulation without inhibiting TDP-43 function (Fig. 6), supporting the notion that nuclear RNA binding sites for TDP-43 can be increased without perturbing TDP-43 function. Synthesizing the recent^27^ and current findings, two key factors enabling TDP-43 functional preservation are: (1) endogenous RNAs as opposed to PS-modified synthetic oligos with increased protein binding affinity,^77^ and (2) a diversity of GU-rich RNA motifs, beyond pure GU-repeats (Fig 5d-g). Diversifying RNA motifs, however, limits or eliminates RBP specificity, as we observed with splicing inhibitors^21^ and 5EU, both of which affect the localization of multiple RBPs, beyond TDP-43.

To date, studies of the RNA-based nuclear retention of TDP-43 have been largely limited to *in vitro* model systems. 5EU has been widely used as an *in vivo* RNA metabolic marker^32,52,61–63^ and mechanistic studies if sufficient doses can be achieved to promote nuclear RNA retention without toxicity. Remarkably, 5EU administered to early postnatal mice was recently shown to persist for up to 2 years, enabling the discovery of a new class of long-lived RNAs in the brain.^78^ However, direct injection of 1 µL of 75 mM 5EU into the cerebellum of adult mice caused Purkinje cell degeneration after 9 days,^64^ demonstrating the risk of neurotoxicity with high doses and the need for dose-finding studies. Identifying the specific molecular target of 5EU that promotes nuclear RNA and RBP accumulation could also facilitate the screening and further development of analogs, with the goal of improving specificity for TDP-43 and limiting toxicity.

Nucleotide and nucleoside analogs have a long history of therapeutic use,^79^ particularly as anticancer agents that compete with physiological nucleosides to induce cytotoxicity^80,81^ and antiviral agents that inhibit viral DNA or RNA polymerases or incorporate into viral nucleic acids causing chain termination.^82^ Notable examples include the adenosine analog, remdesivir, for SARS-CoV-2 ^83^ and the uridine analog sofosbuvir for hepatitis C.^84,85^ There are currently no analogs in therapeutic use for neurodegenerative disease, though preclinical studies recently demonstrated the ability of guanosine/cytidine analogs to mitigate toxicity of the G_4_C_2_ hexanucleotide repeat expansion in a human neuron model of *C9orf72-*ALS/FTD.^86^ Our findings in 5EU-treated cells further suggest that RNA base analogs may have unexpected utility as mechanistic or even therapeutic tools for neurodegeneration.

## EXPERIMENTAL PROCEDURES

### Cell lines

All cells were grown at 37°C in humidified air containing 5% CO_2_. Cell lines were validated by STR profiling (ATCC), verified to be mycoplasma negative (Genlantis), and routinely refreshed from frozen stocks. A monoclonal HeLa-61 cell line (derived from polyclonal ATCC stock)^87^ was maintained OptiMEM (Gibco) with 4% FBS. SHSY5Y cells (ATCC) were maintained in DMEM/F12 (Gibco) with Dasso lab,^88^ and were maintained in DMEM (Gibco) with 10% FBS. Monoclonal HeLa-61-derived cell lines stably expressing V5-tagged wild-type or RRM-mutant TDP-43 were generated by transient transfection followed by puromycin selection and limiting dilution cloning.

To enable inducible CHMP7-FLAG overexpression, gene synthesis (Twist Biosciences) was used to replace TDP43-GFP in pLVX tight puro^89^ with human CHMP7-TEV-3xFLAG flanked by the original BamH1 and NotI sites. Lentiviruses were prepared in HEK293T cells with third-generation packaging vectors (Invitrogen) and transduced into HeLa-61 cells stably expressing tetracycline repressor that was introduced via pLenti CMV rtTA3tx Blast (a gift from Eric Campeau, Addgene plasmid #26429). A single cell-derived line was isolated under blasticidin and puromycin selection and tested for the doxycycline-inducible CHMP7-FLAG expression by immunofluorescence with anti-FLAG (CST).

### Mouse primary neuron culture

Mouse primary cortical neurons were cultured as previously described.^90,91^ Briefly, E16 timed pregnant C57BL/6J female mice (Jackson Laboratory) were sacrificed by cervical dislocation, cortex dissociated, and cells plated at 50,000/well on poly-D-lysine/laminin-coated, optical glass-bottom 96-well plates (CellVis). Neurobasal growth medium was supplemented with B27, Glutamax, and penicillin/streptomycin (Gibco). All animal procedures were approved by the Johns Hopkins Animal Care and Use Committee.

### Human i3-neuron culture

An iPSC line with a doxycycline-inducible neurogenin-2 expression cassette in the AAVS1 safe-harbor locus was provided by the Michael Ward lab.^92^ The iPSCs were maintained in Essential 8 media (ThermoFisher) on Matrigel (Corning) and differentiated into glutamatergic cortical neurons according to the published protocol.^93^

### Immunofluorescence

Paraformaldehyde-fixed cells were rinsed with PBS and blocked and permeabilized with 10% normal goat serum (NGS, Vector Labs) and 0.1% Triton-X-100 in PBS for 15-30 min at room temperature. Primary antibodies (Supplementary Table 1) were applied for 1 h at room temperature or overnight at 4°C in 10% NGS/PBS. After rinsing with PBS, AlexaFluor-labeled secondary antibodies (ThermoFisher) were applied for 1 h at room temperature in 5% NGS/PBS. Finally, cells were rinsed with PBS containing Hoechst 33342 and transferred to 50% glycerol/PBS for imaging.

### RNA fluorescence *in situ* hybridization

Cells were fixed in 10% molecular grade-formaldehyde (Sigma-Aldrich) in PBS for 20 min, permeabilized in 0.1% Triton-X 100/PBS for 10 min, and washed 3 times in 1x PBS and 2 times in 2x SSC (Sigma-Aldrich) for five min. Following prehybridization for 1 h in hybridization buffer at 42°C (Thermo), probe (Supplementary table 2) was applied at 100 nM in hybridization buffer at 42°C overnight. Cells were washed with SSC buffer (2x, 0.5x, and 0.1x in PBS, 20 min each), at 42°C with Hoechst 33342 included in the final wash.

### Click-chemistry labeling

5EU-treated cells were fixed with 4% paraformaldehyde/PBS for 15 min, permeabilized and blocked in 2% BSA/0.1%TX-100 in PBS for 30 min, and labeled with 0.5 μM AF488-Picolyl Azide using the Click-&-Go Cell Reaction Buffer kit according to the manufacturer’s instructions (Click Chemistry Tools).

### *In vitro* RNA transcription

The dsDNA template for *in vitro* RNA transcription was prepared by phosphorylation (using polynucleotide kinase, NEB) and annealing of synthetic DNA oligos (prepared at IDT) containing the (Supplementary table 2). Following the manufacturer’s instructions, the NEB HighScribe T7 RNA Synthesis kit (NEB) was used to synthesize RNAs for 6h at 37°C from equimolar A,G,C,U (NEB) or A,G,C, and 5EU (Jena Biosciences) ribonucleotide mix, in 100ul reactions. Subsequently, the reactions were treated with RNase-free DNase I (NEB) in 1x DNase buffer for 40 min at 37°C to remove the DNA template. RNA was purified using the Monarch RNA Cleanup kit. The RNA concentration was determined with Nanodrop One (Thermo), and the expected size and integrity of the synthetic RNAs was verified by electrophoresis in 2% agarose/TBE gels stained with SYBR Gold (Invitrogen).

### TDP-43 RNA binding assays

Recombinant TDP-43 was generated in *E.coli* and purified as previously described.^41^ TDP-43 intrinsic fluorescence was measured upon titration with increasing concentrations of RNA oligonucleotides and analyzed as previously described.^41^ The assays were conducted in buffer containing 10mM Tris, pH 8.0, 300mM NaCl, 5% glycerol, 5% sucrose, and 1mM TCEP.

### Immunoblots

HeLa cells were treated with or without 100 µM 5EU for 24 h and lysed in RIPA buffer with protease inhibitor cocktail (Roche). After clarification by centrifugation (21,000g, 5 min, 4°C) and protein concentration measurement, samples were boiled in Laemmli sample buffer (Bio-rad) and separated by SDS-PAGE. After transfer to nitrocellulose membranes, immunoblots were blocked with 5% non-fat milk in TBS with 0.05% Tween (pH 7.4, TTBS), incubated overnight at 4°C with primary antibodies, washed with TTBS, and then incubated with horseradish peroxidase-coupled secondary antibodies for 1h at room temperature. ECL images were obtained using ImageQuant LAS400.

### eCLIP

HeLa cells were grown to ∼80% confluence in 15 cm dishes. Following treatment, media was aspirated, cells washed with DPBS, placed on ice, and UV crosslinked at 254-nm (400 mJoules/cm^2^). Cells were scraped and pelleted by centrifugation before snap-freezing in liquid nitrogen. TDP-43 eCLIP experiments and data processing were carried out and analyzed as previously described^94^ utilizing a polyclonal TDP-43 antibody (Proteintech 10782-2-AP). Analyses shown here utilized reproducible peaks (IDR ≤ 0.01, ≥8-fold enriched and p<0.001 in IP versus input) across both replicates as previously described.^94^ For comparison of binding across timepoints, read density (in reads per million) from all four eCLIP conditions was isolated for the region from - 100 to +100nt from the center of each IDR peak in untreated conditions, and divided by the maximum read density across the four conditions. Replicate batches were considered separately as we observed batch-dependent read differences.

For eCLIP k-mer analysis, peaks less than 40nt were extended equally on each size to a minimum size of 40nt, and then each peak was extended in the 5’ direction by 20nt to include motifs lost due to crosslink termination. For fold-enrichment, the background rate for each k-mer was calculated by randomly shuffling the sequence of each peak (Fisher-Yates shuffle) 10 times.

### RNAseq

RNA was extracted from cells using the Zymo Quick-RNA MiniPrep kit according to the manufacturer’s instructions with DNase digestion. Libraries were prepared using Stranded Total RNA Prep Ligation with Ribo-Zero Plus (Illumina). For the RNA decay timecourse, ERCC spike-in controls (Thermo) were added prior to library preparation. Samples were QCed using Qubit measurements for quantitation and fragment analyzer (FA) assay for quality and sizing information. The libraries were pooled together at 4nM based on FA results. The concentration of the library pool was further verified by qPCR before sequencing. Paired-end (2X100) sequencing was done on NovaSeq 6000 S2 200 flow cell.

### RNAseq analysis

Sequencing reads were mapped to the human genome version GRCh38 with the alignment tool STAR v.2.4.2a.^95^ The aligned reads were assembled with PsiCLASS v.1.0.2^96^ to create gene and transcript models, and the assembled transcripts were then assigned to known genes from the human RefSeq reference gene set. Lastly, DESeq2^97^ was used to quantify the expression levels and determine differentially expressed genes. Additional visualizations, including volcano plots and plots of principal coordinate analysis (PCA) components, were produced with custom R scripts.

### Splicing analysis

#### TDP-43 cryptic exons

A custom script was employed to align fastq files using STAR,^95^ extract splice junction counts, and compute percent spliced in (PSI) values for known, TDP-43-dependent cryptic exons in HeLa cells.^98^ A subset of targets were validated by RT-PCR and gel electrophoresis as follows. cDNA synthesis was performed with the high-capacity cDNA reverse transcription kit (ThermoFisher). RT-PCR reactions were performed using Platinum II Hot-Start Green PCR Master Mix (Thermo) and products separated on 2-4% EX e-gels containing SYBR Gold II (Invitrogen) prior to fluorescent imaging with an ImageQuant LAS400 system.

#### Differential splicing

Replicate multivariate analysis of transcript splicing (rMATS)^43^ was used to analyze differences between the ratio of isoforms in canonical alternative splicing events, including exon skipping, mutually exclusive exons, alternative 5’ and 3 splice sites, and intron retention. Events were filtered to include only those with ≥ 10 reads per sample, abs(dPSI)≥ 0.1, and false discovery rate (FDR) ≤ 0.01.

#### Splicing complexity

The tool MntJULiP was used to calculate a per sample measure of splicing complexity as previously described.^44^ MntJULip determines differences in the inclusion levels of genomic region between two consecutive exons and is identified by the pair of coordinates of its two flanking exons. MntJULiP extracts introns and their supporting read counts from spliced RNA-seq read alignments collected from all samples. It then groups introns that share a common endpoint and calculates, for each intron i in the group, a PSI value that represents its contribution to the group’s expression level: PSI(i) = n_i_/(n_1_+n_2_+…+n_k_), where n_i_ is the read count for intron i and k the number of introns. Next, a complexity score s was calculated for the group based on the difference between the PSI value of the primary ‘isoform’, PSI_MAX_, and the group’s PSI average: s = 1-(PSI_MAX_-PSI_avg_). A smaller s value indicates a larger contribution of the primary isoform and therefore lower splicing complexity for the group. Lastly, the scores of all groups were averaged to calculate a per sample measure of splicing complexity.

### RNA decay analysis

HeLa cells were treated ± 100 µM 5EU for 24 h. After washout, 250 nM NVP2 was added for 0, 1, 2, 4, or 8 h prior to RNA extraction and sequencing as above. RNA half-lives were calculated as recently described.^99^ Reads were aligned to a human protein-coding transcriptome reference (Gencode GRCh38.p14) that included the ERCC spike-in sequences, using Salmon.^100^ TPM values at each timepoint were normalized to spike-in (abundance A_(t)_), and further normalized to time 0 (A_0_). Half-lives were then calculated by fitting a nonlinear least squares regression model to the exponential decay model A_(t)_ = A_0_ x e^-kt^. Analyses and plotting were done using R tools such as gplots (heatmap), nlme (regression) and ggplot2 (density, boxplot).

### High content microscopy and automated image analysis

Automated cell imaging was carried out with an ImageXpress Micro Confocal high-content microscope running MetaXpress software (Molecular Devices) as described.^21,27,91^ Briefly, 9-16 non-overlapping fields per well were imaged at 20x (cell lines, i3 neurons) or 40x (primary neurons) in in unbinned, 16-bit images. The background-corrected mean nuclear and cytoplasmic intensities and the nuclear/cytoplasmic (N/C) ratios were calculated using the MetaXpress translocation-enhanced module. Nuclear puncta were quantified using a custom module^27^ to identify puncta from 0.5 to 6 mM in size (1–18 pixels at 20x magnification), with intensity above the local background adjusted for each marker. Hoechst was used to identify the nucleus, and nuclear and cytoplasmic compartments were set several pixels inside and outside the nuclear envelope to avoid edge effects. All image analysis was carried out on raw, unaltered images. Raw data were uniformly filtered to exclude errors of cell identification (probe intensity = 0) and non-physiologic data (e.g. N/C ratios < 0.1 or >100). Where indicated in the figure legends, data were normalized across technical and biological replicates as % untreated controls or percent time 0. The approximate number of cells/well across replicates is also provided.

### Statistical analysis

Statistical analyses and curve fitting were performed in Prism (GraphPad). Differences between groups were analyzed by t-test or ANOVA with post-hoc adjustment for multiple comparisons as detailed in the figure legends. The number of biological replicates (independent experiments and cell passages) was used as the N for all comparisons unless otherwise noted. Dose-response curves were fit by simple linear or non-linear regression.

## Supporting information

Hayes et al Supplement

## DATA AVAILABILITY

All data reported in this paper will be shared by the lead contact (Lindsey Hayes) upon request. This paper does not report original code. RNAseq and eCLIP datasets will be uploaded to the GEO repository and made available at the time of publication.

## ACKNOWLEDGEMENTS

We thank Corina Antonescu from the JHU Computational Biology Consulting Core for the analysis of the RNAseq data, and Svetlana Vidensky, Lyudmila Mamedova, and Fang Yang for their expert technical assistance. This research was supported by the Department of Defense CDMRP AL210092 (LH), the National Institutes of Health R01NS123538 (LH), R35GM144114 (JC); and the Guy McKhann Scholar Award (LH). ELVN is a CPRIT Scholar in Cancer Research (RR200040).

## SUPPLEMENTARY FIGURES

**Figure S1. 5EU time course and source comparison, related to Figure 1.**

**A. Timecourse of 5EU-induced TDP-43 nuclear accumulation.** HeLa cells were treated with 5EU for the indicated amount of time (0-24 h), fixed, and 5EU washed out before transcriptional blockade with ActD for 2 h. The TDP-43 N/C ratio is shown according to the duration of 5EU pretreatment. The mean ± SD of 4 independent replicates is shown, with an average of ∼1900 cells/treatment condition/replicate. NS = not significant, ****p<0.0001 by one-way ANOVA with post-hoc correction for multiple comparisons.

**B-C. Comparison of commercial sources of 5EU.** HeLa cells (B) and mouse primary cortical neurons (C) were pre-treated with 5EU at the indicated doses for 24h before washout and transcriptional blockade with ActD for 2 h. The TDP-43 N/C ratio is shown according to 5EU dose and commercial source (ThermoFisher, Click Chemistry Tools, Carbosynth).

**Figure S2. ‘Click’ labeling of 5EU incorporation across cell types, related to Figure 1.**

**A.** Nuclear AF488-azide intensity in the indicated cell types treated with 5EU for 24 h. Mean ± SD is shown for the number of replicates indicated in the legend.

**B-D.** Representative fluorescent images of AF488-azide-labeled HeLa cells (B), i3-neurons (C), or SHSY5Y cells (D) treated with the indicated concentration of 5EU for 24 h followed by fixation and ‘click’ labeling. The intensity histogram for each image was normalized to the dimmest and brightest pixels of the highest dose i3 neurons. Scale bar = 20 µm.

**Figure S3. 5EU-induced cell cycle blockade, related to Figure 1.**

**A-C.** Total cell count (A) and # metaphase cells (B) in HeLa cells treated with increasing doses of 5EU for 24 h. Sample images of Hoechst staining from untreated vs. 5EU-treated HeLa cells (C) show dose-depending loss of mitotic cells (arrow) and decreasing cell numbers without the appearance of condensed chromatin/dying cells. Scale bar = 50 µm. The mean ± SD for 4 independent replicates is shown.

**D-E**. Cell counts from human i3 neurons (D) and mouse primary cortical neurons (E) treated with increasing doses of 5EU for 24 h show no change in cell number. The mean ± SD for n=3 (D) or n=4 (E) independent replicates is shown. In A-E, NS = not significant, **p<0.01, ****p<0.0001 by one-way ANOVA with post-hoc correction for multiple comparisons.

**Figure S4. 5EU-induced effect on other nuclear RBPs, related to Figure 1.**

**A.** Representative immunofluorescence images of indicated RNA binding proteins (RBPs) in HeLa cells treated with 100 µM 5EU for 24 h prior to washout and transcriptional blockade with 250 nM NVP2 for 2 h. The intensity histogram for each image was spread between the dimmest and brightest pixels. Scale bar = 20 µm.

**B.** N/C ratio for designated RBPs in 5EU-treated HeLa cells with or without NVP2 treatment. Graphs at left show N/C ratio expressed as % untreated cells. Graphs at right compare NVP2-treated vs. untreated (-NVP2) cells to obtain the NVP2-induced shift, corrected for any change in steady-state nuclear RBP levels. The mean ± SD of 2 independent replicates is shown, from an average of ∼2000 cells/treatment condition/replicate.

**C-D**. Representative immunoblots (C) for indicated RBPs in untreated vs. 5EU-treated HeLa cells (100 µM, 24 h) and protein abundance (D) in n=5 independent replicates normalized to GAPDH. NS = not significant, one-way ANOVA with post-hoc correction for multiple comparisons.

**Figure S5. TDP-43 affinity testing via fluorescence-based binding assay, related to Figure 3.**

**A-C**. The apparent dissociation constant (*K_D,app_*) of TDP-43 for *in vitro-*transcribed ‘AUG12’ without (A) or with (B) 5EU was measured by a fluorescence-based assay that monitors the expected decrease in the intrinsic fluorescence of recombinant TDP-43 upon RNA binding. Fluorescence with and without oligo (*F/F_0_)* was measured for increasing RNA concentrations, and the best fit values for *K_D,app_* were calculated as previously described from N=4 binding curves per oligo.^41^

**Figure S6. Time course of 5EU-induced changes in RNA abundance/localization, related to Figure 4.**

**A.** Representative 488-azide ‘click’ labeling of 5EU-containing RNA in HeLa cells treated with 5EU for 0, 1, 6, and 24 h. The intensity histogram in the top row of images was normalized to the brightest condition (24 h). In the lower set of images, the intensity histogram for each image was spread between the dimmest and brightest pixels. A pseudo-color linear LUT was applied to facilitate visualization of the cytoplasmic signal (arrows). Scale bar = 20 µm.

**B.** N/C ratio (upper) and nuclear/cytoplasmic intensities (lower) for ‘click’ label signal over time. Mean ± SD from N∼5000 cells/treatment condition.

**C-D**. Poly(A) (C) and (GU)6-FISH (D) in HeLa cells treated with 100 µM 5EU for 1, 6, or 24 h. Mean ± SD N/C ratio (upper) and nuclear/cytoplasmic intensities (lower) are shown from N∼4000 cells/treatment condition.

**Figure S7. Extended eCLIP data, related to Figure 5.**

**A.** 200nt windows centered on each TDP-43 eCLIP peak in untreated vs. 5EU ± NVP2-treated cells. Each row indicates the normalized read density. Data are from replicate 2, which was normalized separately from replicate 1 (Fig. 5B) to control for batch effects.

**B-C.** Bars indicate the log_2_(fold-enrichment) for indicated 6-mers from significant, reproducible TDP-43 eCLIP peaks from the indicated conditions.

**Figure S8. 5EU-induced changes in nuclear bodies, related to Figure 8.**

**A, C, E**. Representative immunofluorescence staining for Cajal bodies (A, coilin), nuclear speckles (C, sc35), and nucleoli (E, fibrillarin) in untreated vs. 5EU-treated HeLa cells. Scale bar = 20 µm.

**B, D, F.** Quantification of nuclear puncta, total puncta area, and mean puncta size. The mean ± SD from 3 independent replicates is shown, from an average of ∼2100 cells/treatment condition/replicate. NS= not significant, *p<0.05, **p<0.01, ***p<0.001, ****p<0.0001 by one-way ANOVA with post-hoc correction for multiple comparisons. All comparisons vs. untreated cells.

**Figure S9. Uncropped gels, related to Figure 6b and supplementary Figure 4c.**

