## Supplementary material for "5-ethynyluridine perturbs nuclear RNA metabolism to promote the nuclear accumulation of TDP-43 and other RNA binding proteins": Hayes et al Supplement

### Suppl Figure 1

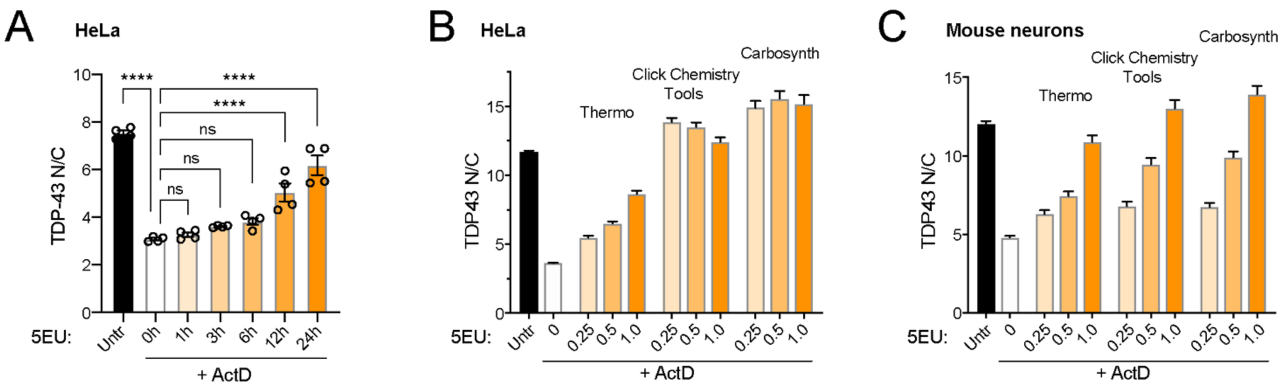


**Figure S1. 5EU time course and source comparison, related to Figure 1**.

**A.** Timecourse of 5EU-induced TDP-43 nuclear accumulation. HeLa cells were treated with 5EU for the indicated amount of time (0-24 h), fixed, and 5EU washed out before transcriptional blockade with ActD for 2 h. The TDP-43 N/C ratio is shown according to the duration of 5EU pretreatment. The mean ± SD of 4 independent replicates is shown, with an average of ~1900 cells/treatment condition/replicate. NS = not significant, ****p<0.0001 by one-way ANOVA with post-hoc correction for multiple comparisons.

**B-C.** Comparison of commercial sources of 5EU. HeLa cells (B) and mouse primary cortical neurons (C) were pre-treated with 5EU at the indicated doses for 24h before washout and transcriptional blockade with ActD for 2 h. The TDP-43 N/C ratio is shown according to 5EU dose and commercial source (ThermoFisher, Click Chemistry Tools, Carbosynth).

### Suppl Figure 2

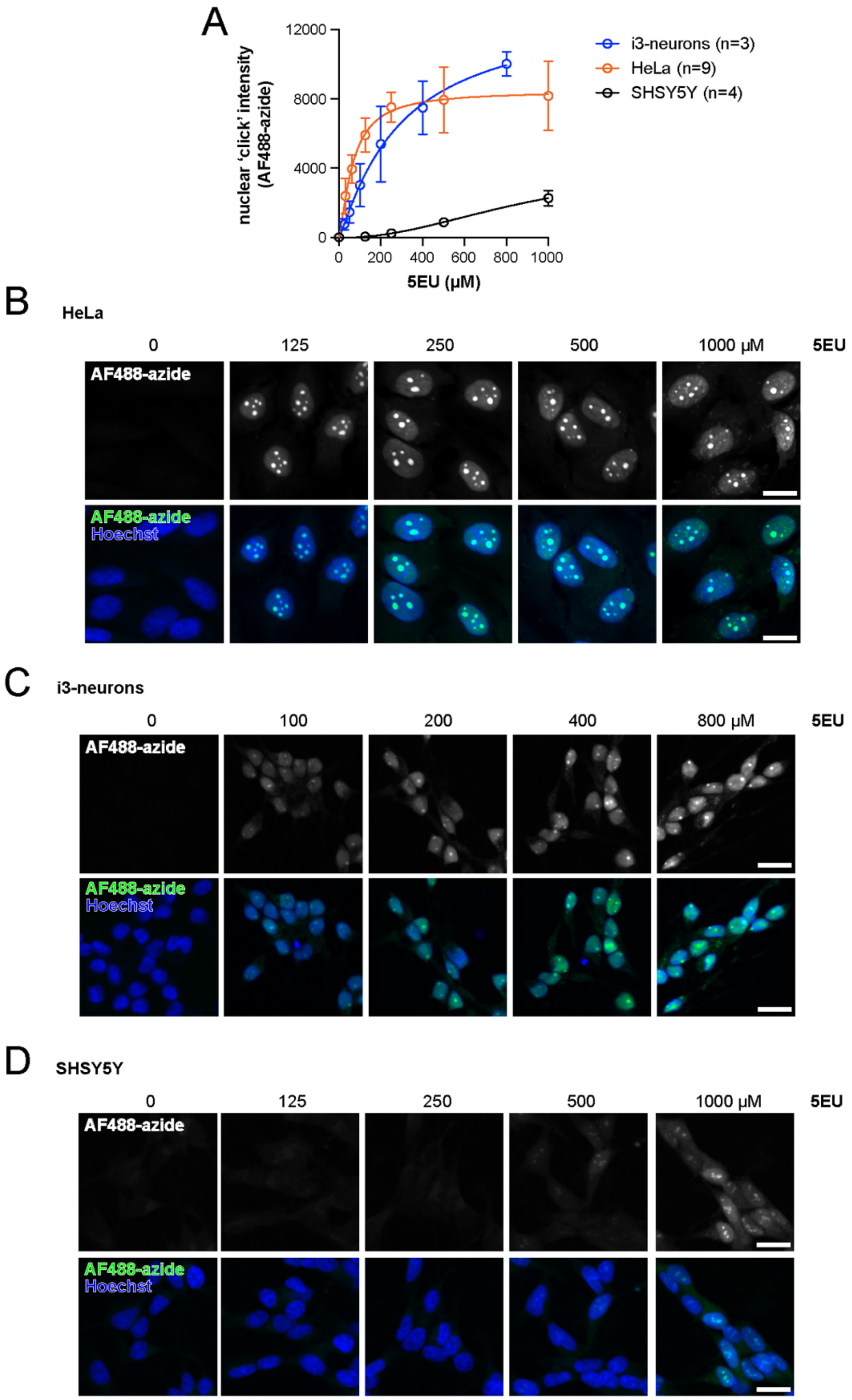


**Figure S2. 'Click' labeling of 5EU incorporation across cell types, related to Figure 1.**

**A.** Nuclear AF488-azide intensity in the indicated cell types treated with 5EU for 24 h. Mean ± SD is shown for the number of replicates indicated in the legend.

**B-D.** Representative fluorescent images of AF488-azide-labeled HeLa cells (B), i3-neurons (C), or SHSY5Y cells (D) treated with the indicated concentration of 5EU for 24 h followed by fixation and 'click' labeling. The intensity histogram for each image was normalized to the dimmest and brightest pixels of the highest dose i3 neurons. Scale bar = 20 µm.

Suppl Figure 3

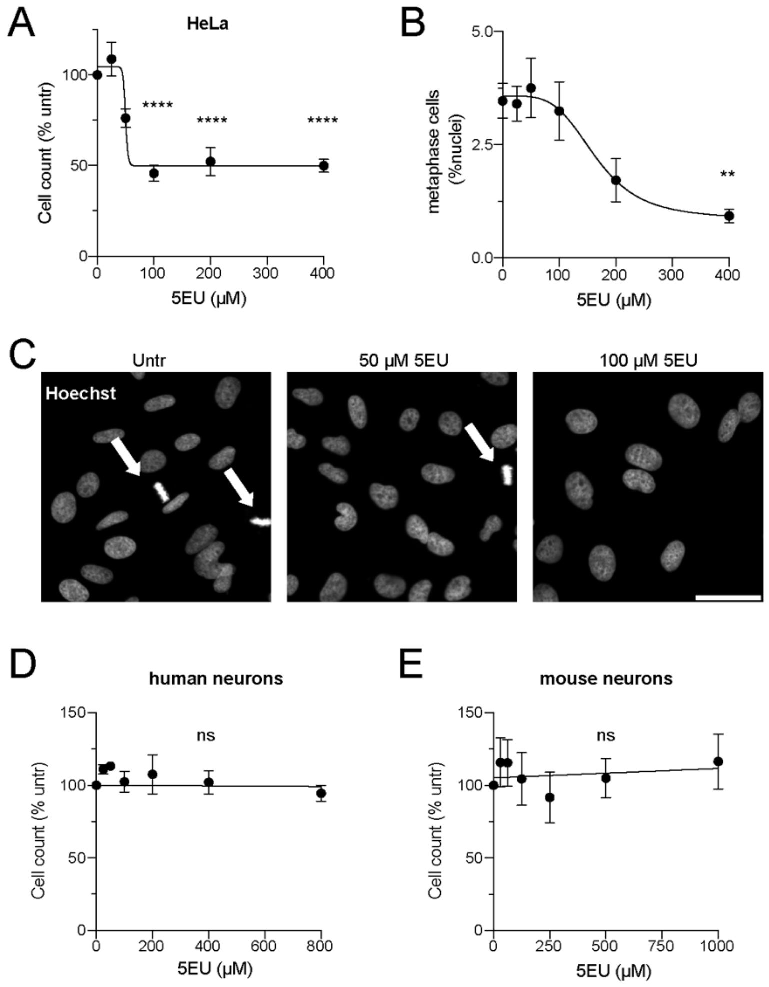


**Figure S3. 5EU-induced cell cycle blockade, related to Figure 1**.

**A-C.** Total cell count (A) and # metaphase cells (B) in HeLa cells treated with increasing doses of 5EU for 24 h. Sample images of Hoechst staining from untreated vs. 5EU-treated HeLa cells (C) show dose-depending loss of mitotic cells (arrow) and decreasing cell numbers without the appearance of condensed chromatin/dying cells. Scale bar = 50 µm. The mean ± SD for 4 independent replicates is shown.

**D-E**. Cell counts from human i3 neurons (D) and mouse primary cortical neurons (E) treated with increasing doses of 5EU for 24 h show no change in cell number. The mean ± SD for n=3 (D) or n=4 (E) independent replicates is shown.

In A-E, NS = not significant, **$p<0.01$, ****$p<0.0001$ by one-way ANOVA with post-hoc correction for multiple comparisons.

#### Suppl Figure 4

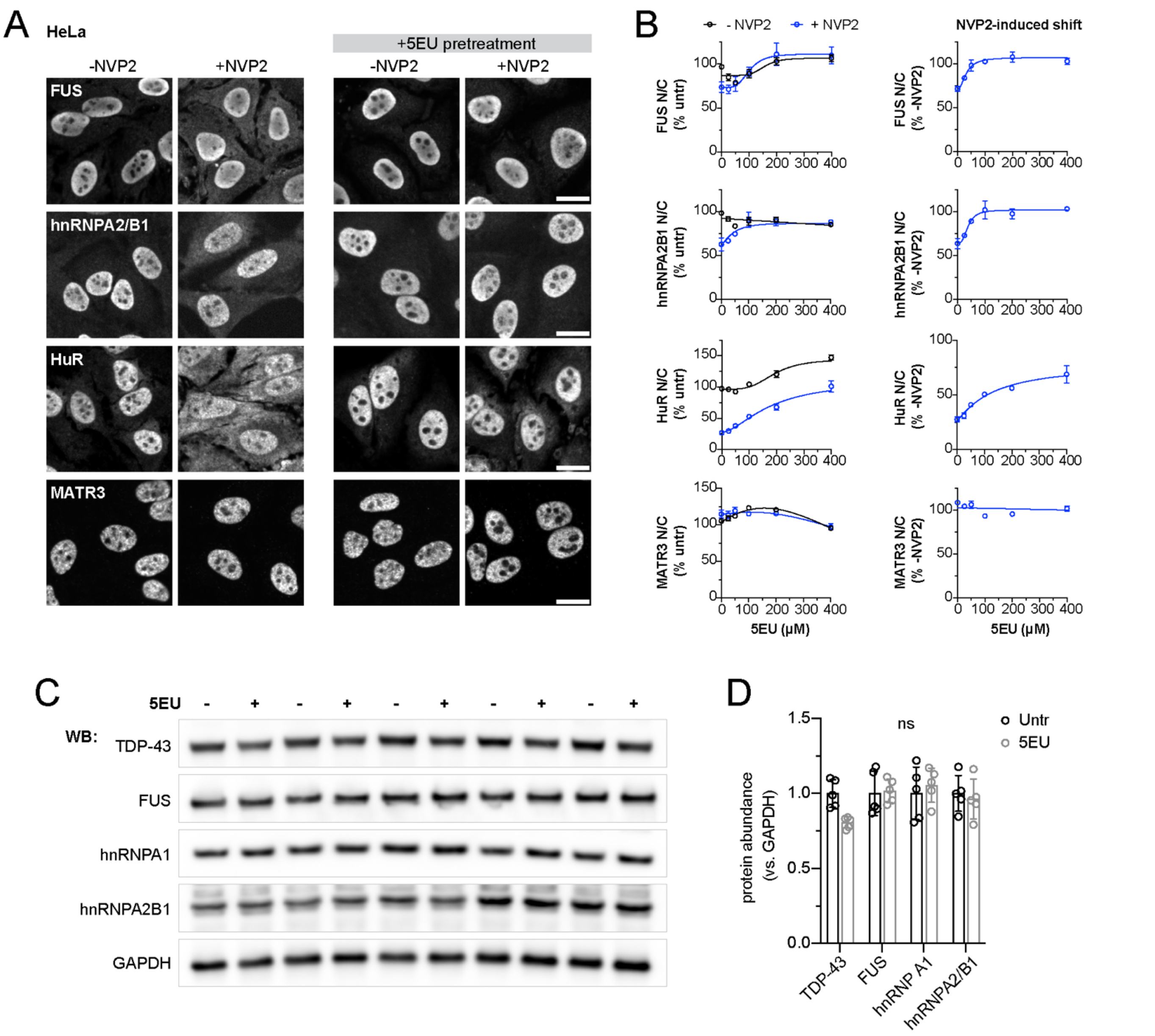


**Figure S4. 5EU-induced effect on other RBPs, related to Figure 1**.

**A.** Representative immunofluorescence images of indicated RNA binding proteins (RBPs) in HeLa cells treated with 100 µM 5EU for 24 h prior to washout and transcriptional blockade with 250 nM NVP2 for 2 h. The intensity histogram for each image was spread between the dimmest and brightest pixels. Scale bar = 20 µm.

**B.** N/C ratio for designated RBPs in 5EU-treated HeLa cells with or without NVP2 treatment. Graphs at left show N/C ratio expressed as % untreated cells. Graphs at right compare NVP2-treated vs. untreated (-NVP2) cells to obtain the NVP2-induced shift, corrected for any change in steady-state nuclear RBP levels. The mean ± SD of 2 independent replicates is shown, from an average of ~2000 cells/treatment condition/replicate.

**C-D**. Representative immunoblots (C) for indicated RBPs in untreated vs. 5EU-treated HeLa cells (100 µM, 24 h) and protein abundance (D) in n=5 independent replicates normalized to GAPDH. NS = not significant, one-way ANOVA with post-hoc correction for multiple comparisons.

Suppl Figure 5

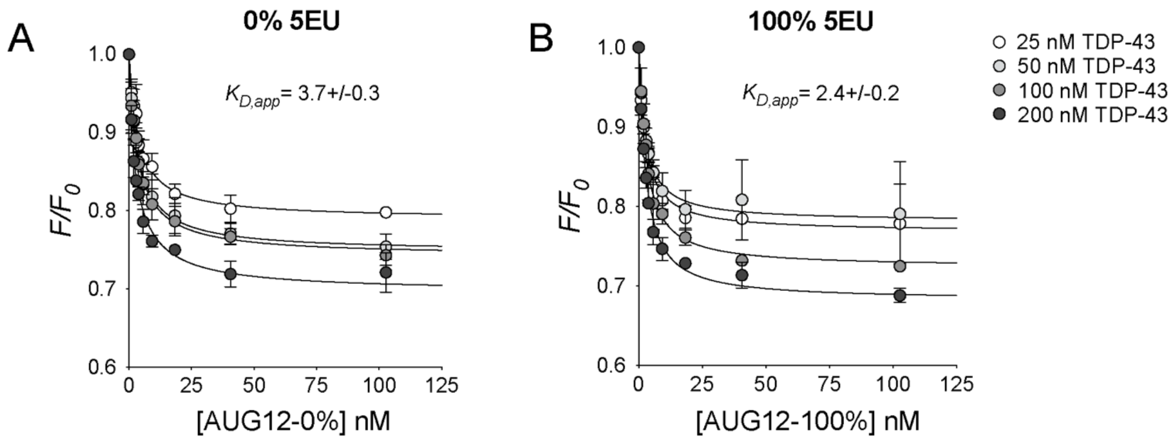


**Figure S5. TDP-43 affinity testing via fluorescence-based binding assay, related to Figure 3**.
**A-C**. The apparent dissociation constant ($K_{D,app}$) of TDP-43 for *in vitro*-transcribed 'AUG12' without (A) or with (B) 5EU was measured by a fluorescence-based assay that monitors the expected decrease in the intrinsic fluorescence of recombinant TDP-43 upon RNA binding. Fluorescence with and without oligo ($F/F_0$) was measured for increasing RNA concentrations, and the best fit values for $K_{D,app}$ were calculated as previously described from N=4 binding curves per oligo.[41]

### Suppl Figure 6

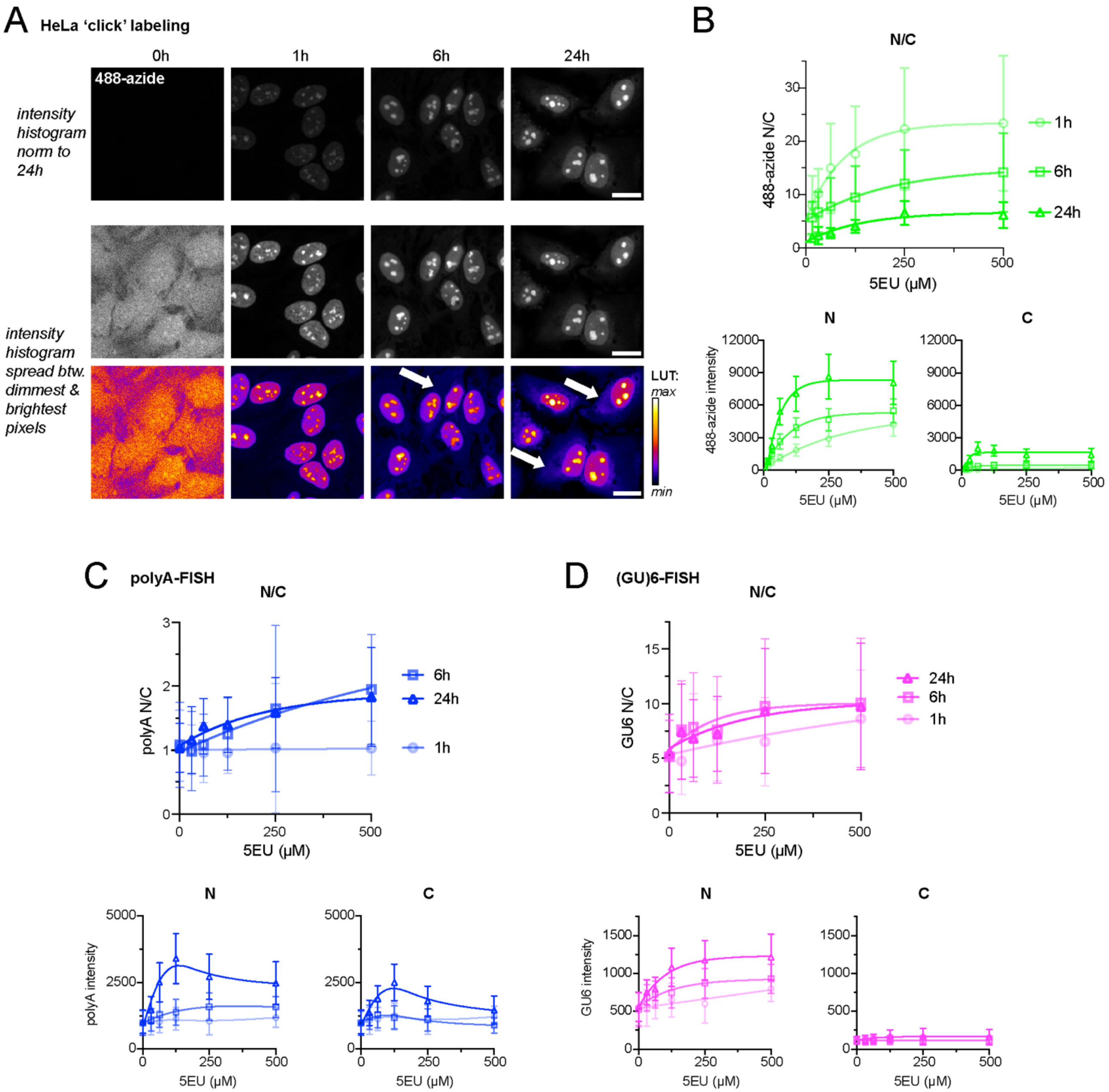


**Figure S6. Time course of 5EU-induced changes in RNA abundance/localization, related to Figure 4**.

**A.** Representative 488-azide 'click' labeling of 5EU-containing RNA in HeLa cells treated with 5EU for 0, 1, 6, and 24 h. The intensity histogram in the top row of images was normalized to the brightest condition (24 h). In the lower set of images, the intensity histogram for each image was spread between the dimmest and brightest pixels. A pseudo-color linear LUT was applied to facilitate visualization of the cytoplasmic signal (arrows). Scale bar = 20 μm.

**B.** N/C ratio (upper) and nuclear/cytoplasmic intensities (lower) for 'click' label signal over time. Mean ± SD from N~5000 cells/treatment condition.

**C-D**. Poly(A) (C) and (GU)6-FISH (D) in HeLa cells treated with 100 μM 5EU for 1, 6, or 24 h. Mean ± SD N/C ratio (upper) and nuclear/cytoplasmic intensities (lower) are shown from N~4000 cells/treatment condition.

### Suppl Figure 7

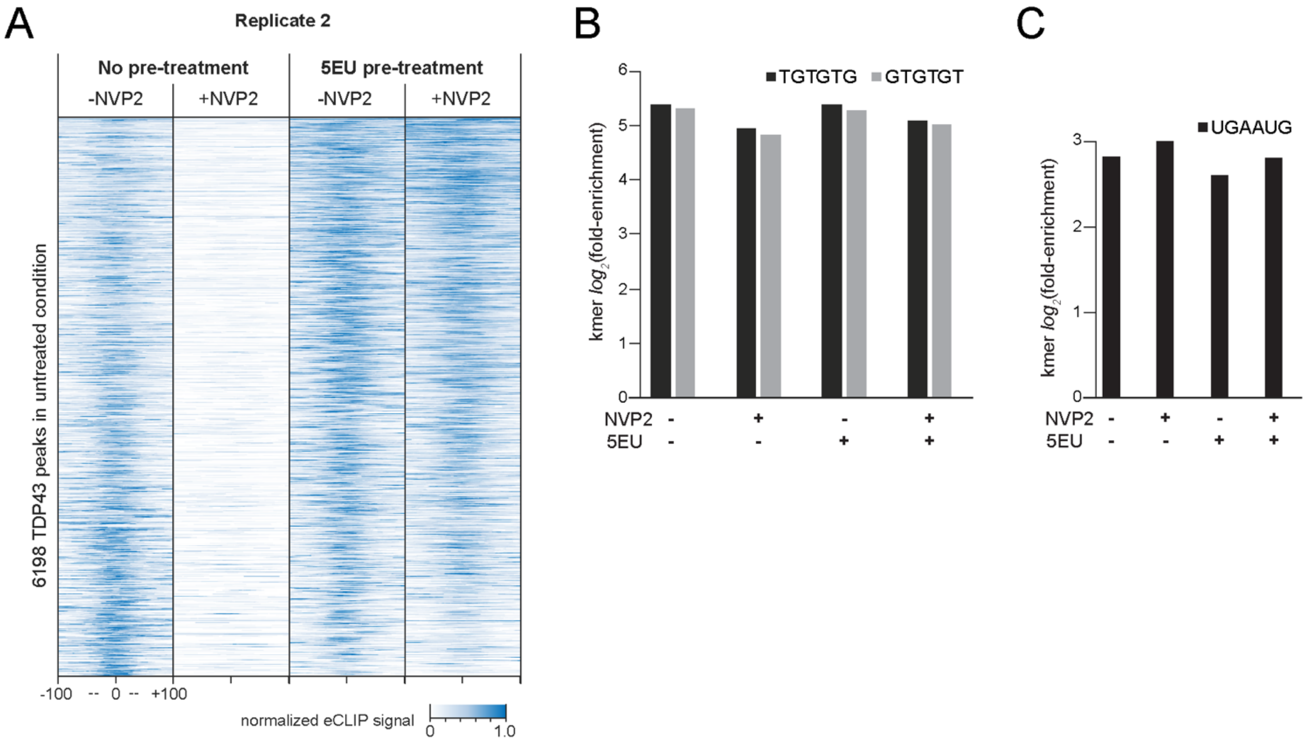


**Figure S7. Extended eCLIP data, related to Figure 5**.

**A.** 200nt windows centered on each TDP-43 eCLIP peak in untreated vs. 5EU ± NVP2-treated cells. Each row indicates the normalized read density. Data are from replicate 2, which was normalized separately from replicate 1 (Fig. 5b) to control for batch effects.

**B-C.** Bars indicate the $\log_2$(fold-enrichment) for indicated 6-mers from significant, reproducible TDP-43 eCLIP peaks from the indicated conditions.

Suppl Figure 8

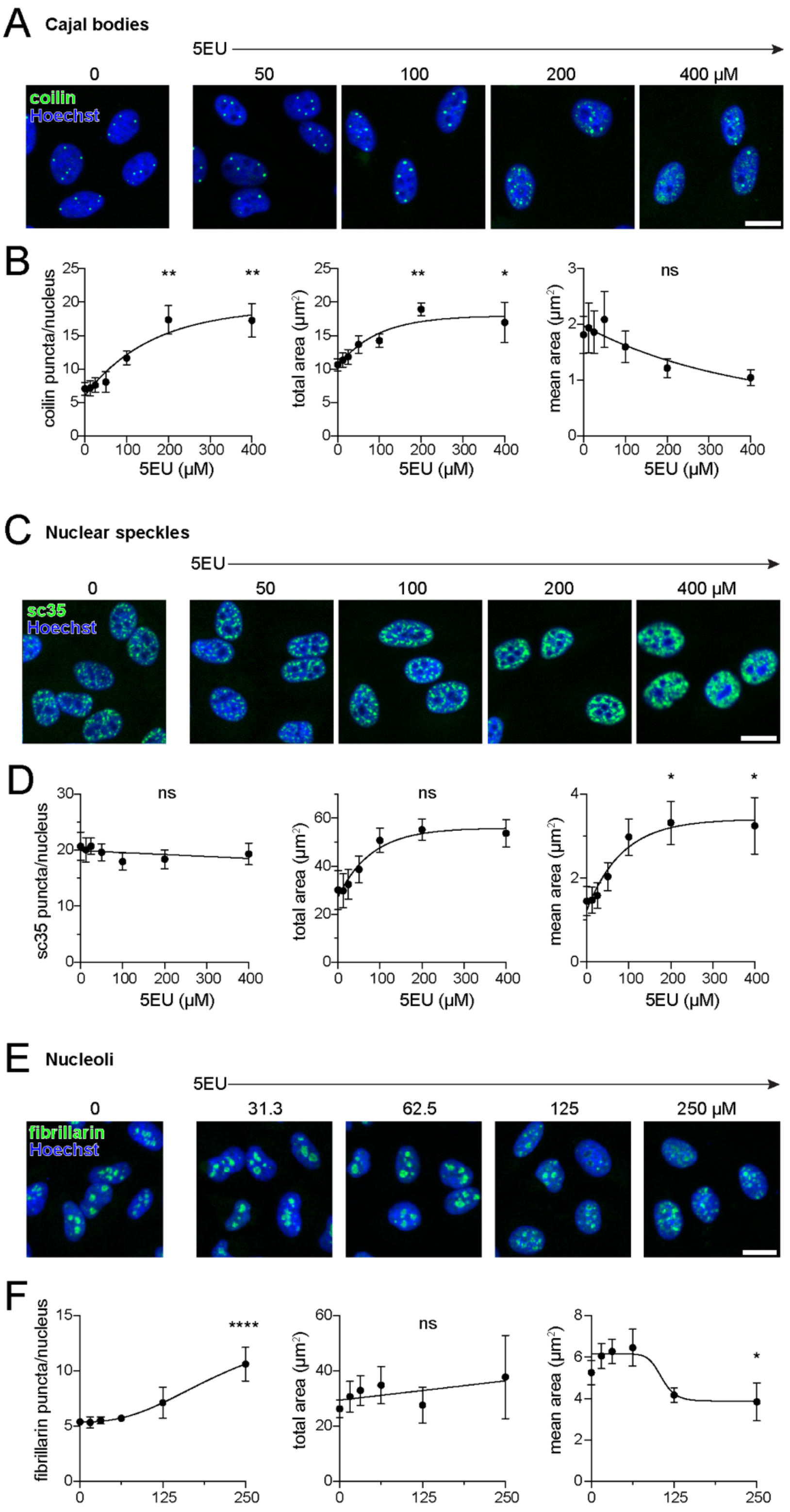


**Figure S8. 5EU-induced changes in nuclear bodies, related to Figure 8**.

**A, C, E**. Representative immunofluorescence staining for Cajal bodies (A, coilin), nuclear speckles (C, sc35), and nucleoli (E, fibrillarin) in untreated vs. 5EU-treated HeLa cells. Scale bar = 20 μm.

**B, D, F.** Quantification of nuclear puncta, total puncta area, and mean puncta size. The mean ± SD from 3 independent replicates is shown, from an average of ~2100 cells/treatment condition/replicate. NS= not significant, *p<0.05, **p<0.01, ***p<0.001, ****p<0.0001 by one-way ANOVA with post-hoc correction for multiple comparisons. All comparisons vs. untreated cells.

### Suppl Figure 9

**Fig 3F**

**EPB41L4A**

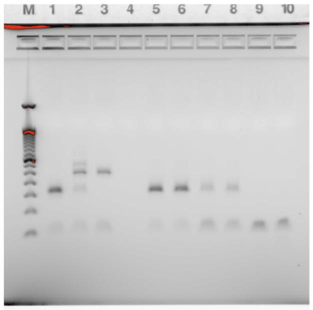


**POLDIP3**

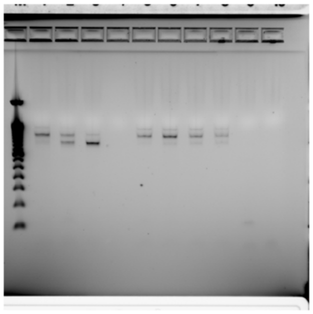

**ARHGAP32**

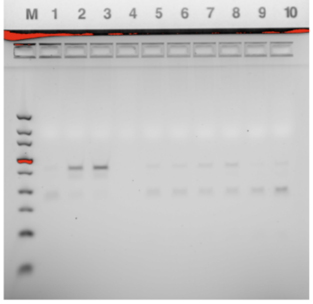


**GAPDH**

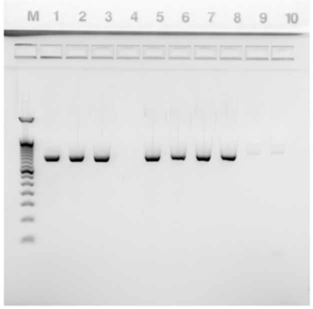


**Suppl Fig 2**

**TDP-43 (43 kD)**

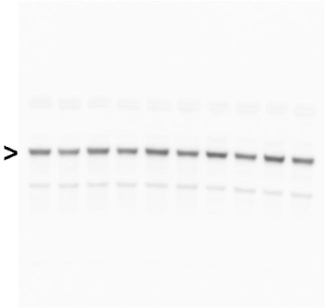

**FUS (75 kD)**

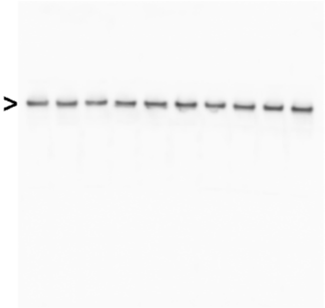

**HNRNPA1 (35 kD)**

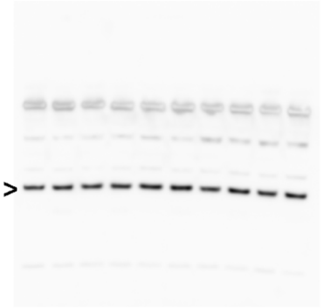

**HNRNPA2B1 (37 kD)**

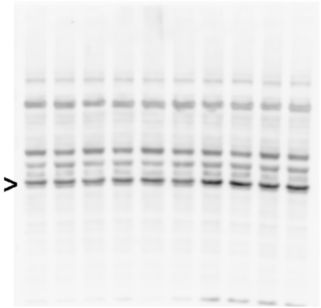

**GAPDH (36 kD)**

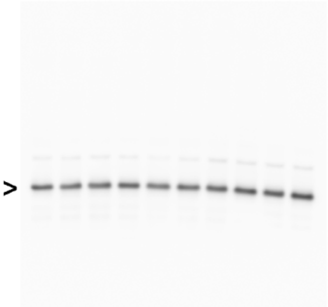

**Figure S9. Uncropped gels, related to Figure 6b and supplementary Figure 4c**

**SUPPLEMENTARY TABLES**

**Supplementary table 1. Primary antibodies**

| **Antibody** | **Source** | **Identifier** |
|---|---|---|
| Rabbit anti-TDP43 polyclonal | Proteintech | Cat# 10782-2-AP<br>RRID: AB_615042 |
| Mouse anti-hnRNP A1 monoclonal (IF) | Santa Cruz Biotechnology | Cat# sc-32301;<br>RRID: AB_627729 |
| Rabbit anti-hnRNP A1 polyclonal (WB) | Proteintech | Cat# 11176-1-AP<br>RRID: AB_2117177 |
| Mouse anti-hnRNP A2/B1 monoclonal | Santa Cruz Biotechnology | Cat# sc-53531;<br>RRID: AB_2248245 |
| Mouse anti-hnRNP K monoclonal | Santa Cruz Biotechnology | Cat# sc-28380;<br>RRID: AB_627734 |
| Rabbit anti-FUS polyclonal | Bethyl | Cat# A300-302A;<br>RRID: AB_309445 |
| Rabbit anti-Matrin 3 monoclonal | Abcam | Cat# ab151714;<br>RRID: AB_2491618 |
| Rabbit anti-ELAVL1/HuR polyclonal | Proteintech | Cat# 11910-1-AP;<br>RRID: AB_11182183 |
| Mouse anti-sc35 monoclonal | Sigma-Aldrich | Cat# S4045;<br>RRID: AB_477511 |
| Rabbit anti-fibrillarin polyclonal | Abcam | Cat# ab5821;<br>RRID: AB_2105785 |
| Mouse anti-coilin monoclonal | Abcam | Cat# ab87913;<br>RRID: AB_10860831 |
| Rabbit anti-GAPDH monoclonal | Cell Signaling Technology | Cat# 2118;<br>RRID:AB_561053 |
| Rabbit anti-V5 monoclonal | Cell Signaling Technology | Cat# 13202;<br>RRID: AB_2687461 |
| Mouse anti-FLAG monoclonal | Cell Signaling Technology | Cat#8146S;<br>RRID: AB_10950495 |

**Supplementary table 2. Oligonucleotide, primer, and probe sequences**

| **Target** | **Sequence**<br>r: RNA base | **Source** |
|---|---|---|
| RNA FISH probes | | |
| Oligo-dT(45)-FAM | T(45) | IDT |
| CA6-Cy5 | rUrCrArCrArCrArCrArCrArCrA/3Cy5Sp/ | IDT |
| DNA oligos for 'AUG12' *in vitro* transcription | | |
| OK1031 | GGCAAAGAACCGTAATACGACTCACTATAGGAAAAAAAAA<br>GTGTGAATGAATAAAAAAAAAAAAAAAAAA | IDT |
| OK1036 | TTTTTTTTTTTTTTTTTTTATTCATTCACACTTTTTTTTTCCT<br>ATAGTGAGTCGTATTACGGTTCTTTGCC | IDT |
| Primers | | |
| EPB41L4A-F | GGACCTCCATATACTTTGTATTTTGGT | Tan *et al*., 2016 |
| EPB41L4A-R | AGCTGAGCAGCAGTGTTGAC | Tan *et al*., 2016 |
| ARHGAP32-F | GAGGGTGTTTGGTTGTGACC | Tan *et al*., 2016 |
| ARHGAP32-R | TTCAGGGAGATGGTTTCCAG | Tan *et al*., 2016 |
| POLDIP3-F | GCTTAATGCCAGACCGGGAGTTG | Ferguson *et al.,* 2019 |
| POLDIP3-R | TCATCTTCATCCAGGTCATATAAATT | Ferguson *et al.,* 2019 |
| GAPDH-F | ACCACAGTCCATGCCATCAC | IDT |
| GAPDH-R | TCCACCACCCTGTTGCTGTA | IDT |
